# The Concise Language Paradigm in left and right hemisphere stroke: a longitudinal source-localized EEG study

**DOI:** 10.64898/2026.10.22.759366

**Authors:** Natascha M. Roos, Robert Oostenveld, Maria Carla Piastra, Anil M. Tuladhar, Marina B. Ruiter, Roy P.C. Kessels, Britta U. Westner, Vitória Piai

**Affiliations:** Department of Neuropsychology and Rehabilitation Psychology, Donders Centre for Cognition, Donders Institute for Brain, Cognition, and Behaviour, Radboud University, Nijmegen, the Netherlands; Donders Centre for Cognitive Neuroimaging, Donders Institute for Brain, Cognition, and Behaviour, Radboud University, Nijmegen, the Netherlands; NatMEG, Karolinska Institute, Stockholm, Sweden; Clinical Neurophysiology, Technical Medical Centre, Faculty of Science and Technology, University of Twente, Enschede, Netherlands; Department of Neurology, Donders Institute for Brain, Cognition, and Behaviour, Radboud University Medical Center, Nijmegen, the Netherlands; Centre for Language Studies, Radboud University, Nijmegen, The Netherlands; Vincent van Gogh Institute for Psychiatry, d’n Herk 90, 5803 DN Venray The Netherlands; Klimmendaal Rehabilitation Center, Heijenoordseweg 5, 6813 GG, Arnhem, The Netherlands; Radboud University Medical Center, Radboudumc Alzheimer Center, Geert Grooteplein Zuid 10, 6525 GA Nijmegen, The Netherlands; Department of Medical Neuroscience, Donders Institute for Brain, Cognition, and Behaviour, Radboud University Medical Center, Nijmegen, the Netherlands

## Abstract

Language recovery after stroke can be supported by neuroplasticity, but its underlying mechanisms remain poorly understood. Most previous studies have either focused on language comprehension or production and/or relied on hemodynamic measures, with limited complementary electrophysiological evidence. Here, we combined a paradigm assessing comprehension and production with individualized electroencephalography (EEG) source reconstruction to investigate post-stroke language function and changes in hemispheric dominance over time.

We tested 31 participants: 13 stroke-free controls, 8 individuals with left hemisphere (LH), and 10 with right hemisphere (RH) stroke in the middle cerebral artery territory. Participants with stroke were assessed approximately two and seven months post-stroke. We employed the Concise Language Paradigm (CLaP), which combines auditory sentence comprehension with picture naming, in addition to standardized neuropsychological tests. EEG data were analyzed using event-related potentials and time-frequency representations, followed by source reconstruction based on individual MRI-derived forward models. Laterality indices were calculated to assess changes in hemispheric dominance between sessions.

Behaviorally, the LH group showed low performance at two months followed by overall improvement, whereas the RH group mainly improved in visuospatial memory and functional communication. EEG activity was significantly modulated by task condition across comprehension and production contrasts. Source-level results in controls were consistent with established language-related networks, serving as regions of interest for the stroke analyses. In the stroke groups, hemispheric dominance differed between LH and RH stroke participants. Notably, several LH participants shifted from left-to right-hemisphere dominance between sessions, whereas those initially showing right-hemisphere dominance remained stable.

Our study provides evidence on comprehension *and* production performance after stroke using a single paradigm. These findings demonstrate dynamic changes in hemispheric dominance during post-stroke recovery, ultimately highlighting the value of individualized EEG source reconstruction to investigate post-stroke plasticity.

## 1 Introduction

Left hemisphere (LH) stroke is a common cause of language and communication impairments, affecting everyday life (Berthier, 2005; Bullier et al., 2020; Hilari et al., 2003). Such impairments tend to be most severe in the first weeks after a stroke, but can recover over time to varying extent (Schellekens et al., 2024; Wilson et al., 2023). Spontaneous recovery^1^ of language function, as induced by neuroplasticity, mostly occurs within the first three months after stroke (Berthier, 2005; Kauhanen et al., 2000), and can be categorized into three phases (Hillis et al., 2006; Kiran, 2012). In the early sub-acute phase (first weeks), the restoration of cerebral blood flow plays a critical role (Hillis & Heidler, 2002). During the late sub-acute phase (approx. 3-6 months), improvement of language function has been associated with the resolution of diaschisis, a functional disruption in distant brain areas connected to the lesioned area (Cappa et al., 1997). Finally, in the chronic phase (> 9 months), further spontaneous improvement can occur, with an ongoing discussion regarding the recruitment of additional brain areas.

It still remains subject to debate whether a (temporary) involvement of right hemisphere (RH) areas supports improvement of language functions or reflects interhemispheric disinhibition, maladaptive for language recovery (Kiran, 2012; Price & Crinion, 2005; Saur et al., 2006). However, most literature suggests that recovery depends on lesion size and location (Schneck et al., 2026; Wilson et al., 2023), and that the best behavioral outcome is linked to pre-stroke organization of brain functions (Coleman et al., 2017; Cramer, 2008). Finally, neuroplasticity usually declines with age, which is why young stroke survivors not only tend to recover faster, but also have higher chances of complete cognitive recovery (Roberts et al., 2026; Watila & Balarabe, 2015; but see Chupina et al., 2022).

Previous approaches to investigate language recovery after stroke majorly differ in the extent of data collected, for example by focusing either on behavioural or neural measures including only one session or not including a control group (Wilson & Schneck, 2021). In addition, studies typically focus on a LH stroke group and therefore cannot discriminate between overall effects of stroke (in either hemisphere) versus effects of LH versus RH stroke specifically. In other words, the inclusion of a RH stroke group allows to determine whether hemispheric stroke location differentially affects specific outcomes.

Most previous studies focusing on neuroplasticity used only one task to assess language, typically yielding one outcome measure of either comprehension or production. This is potentially problematic because language functions can dissociate following stroke: impairments and recovery in one domain may not necessarily be reflected in the other (El Hachioui et al., 2013). Consequently, relying on a single language measure may provide only a partial picture of post-stroke language functioning and its neural correlates. To address this, we have designed the Concise Language Paradigm (CLaP), a sentence-embedded picture-naming task that comprises measures of language comprehension and production within one trial (Roos et al., 2024). The different contrasts of the CLaP evaluate auditory sentence comprehension during the sentence interval, contextually guided word retrieval during the pre-picture interval, and visually guided word retrieval during the picture interval. Especially the context effect between contextually and visually guided word retrieval has been demonstrated to repeatedly involve the left temporo-parietal cortex using electrophysiology as well as fMRI (Klaus et al., 2020; Piai et al., 2014, 2015, 2017, 2018; Roos et al., 2023; Roos & Piai, 2020). Although combining comprehension and production within a single task introduces the possibility of task impurity (e.g., impairments in one domain could affect performance in the other), the ability to assess both functions within the same paradigm offers the important opportunity of a more integrated characterization of post-stroke language function and neuroplasticity. Thus, by evaluating processes of language comprehension as well as production, the CLaP provides a suitable framework to investigate language recovery after stroke, in light of previously established group-level results in a stroke-free sample (Roos et al., 2024).

In addition to the choice of how and which language functions are assessed, the measures used to investigate the underlying neural activity also warrant careful consideration. A recent systematic review and meta-analysis of functional imaging studies on post-stroke language plasticity illustrates a lack of compelling evidence and methodological consistency in the literature (Wilson & Schneck, 2021). Most previous studies have relied on hemodynamic measures, such as fMRI, while complementary electrophysiological evidence remains scarce. However, accumulating evidence based on only one type of measurement has important limitations (for discussion Piai & Francken, 2026), particularly in the context of brain damage. Both hemodynamic and electrophysiological signals can be affected by stroke and yield invalid or misleading results (Piai et al., 2024; Piastra et al., 2022; Veldsman et al., 2015), an issue that is not always acknowledged. As an indirect measure of neural activity, fMRI may be suboptimal for estimating neural activity based on hemodynamic responses after stroke due to changes in neurovascular coupling (Lindauer et al., 2010; Veldsman et al., 2015). Moreover, the poor temporal resolution of the hemodynamic response may introduce interpretation issues when it comes to processes that are highly dynamic, such as those involved in language (e.g., Hauk, 2016).

The caveat of relying on cerebrovascular hemodynamics to infer neural activity following stroke has been highlighted in a combined MEG and fMRI study, testing stroke-free controls as well as LH and RH stroke participants (Rossini et al., 2004). The authors used electric nerve stimulation, which reliably elicited symmetric bilateral responses in controls with MEG as well as fMRI. For the stroke participants, MEG yielded clearly detectable responses in both hemispheres, regardless of stroke location, whereas the detection of the corresponding BOLD signal mostly failed. This absence was not determined by the location or size of the lesion, as the BOLD signal was absent in both hemispheres in more than half of the stroke participants. In the absence of the MEG measurements, one might have interpreted these abnormal BOLD responses to reflect neuroplasticity-induced changes, especially in cases where the signal was only lacking in the affected hemisphere. This demonstration of how hemodynamics can become uncoupled from neural activity following stroke stresses the importance of incorporating complementary evidence from electrophysiology. In comparison to fMRI, electrophysiology affords capturing time-dynamic changes in neural activity during specific time windows, affording more specific interpretation in relation to task performance.

Electrophysiological measures, in contrast to fMRI, provide a direct measure of neural activity with millisecond resolution, and can thus provide valuable insights into fast-paced neural processes during language use after stroke. However, EEG scalp-level results are affected by stroke lesions, which asymmetrically impact the spatial distribution and conductivity of brain tissue, affecting the flow of electrical currents through the brain. However, these issues can be taken into account by employing source reconstruction techniques with a high level of methodological sophistication (Piai et al., 2024; Piastra et al., 2022). This requires participant-specific forward models based on individual structural MRI scans and detailed modelling of the stroke lesion in the brain. However, this approach is very rarely employed. For example, a recent review of longitudinal electrophysiology studies on post-stroke aphasia between 1950 and 2022 comprised only 34 studies (Arheix-Parras et al., 2023). Only five of these included source reconstruction techniques, and none of them used such individualized forward models. Another recent review criticised the fact that most post-stroke aphasia studies do not address the impact of stroke lesions on scalp and source-level findings (Lu et al., 2024; but see Chupina et al., 2022; Piai et al., 2017 for examples where this was done).

### 1.1 Present study

Given the methodological limitations of current language tasks and of the neuroimaging techniques discussed above, the focus of the present study was to gather evidence on combined comprehension *and* production performance after stroke using state-of-the-art electrophysiology analyses at the scalp and source level. We therefore applied recent methodological improvements in the field of biophysical modelling to the study of language functioning. Adopting these recent developments would help advance the understanding of neuroplasticity during recovery of language after stroke. Our aim was to establish if the time-specific cortical responses associated with comprehension and production would be observed in the LH and RH stroke groups, and if these responses would change (spatially) from the late sub-acute to the chronic phase. Given the direct nature of the electrophysiological responses in terms of the underlying neural activity, (changes in) these cortical responses would afford a degree of interpretability complementary to what is currently available in the literature.

Using a longitudinal design with behavioral, EEG, and MRI data, we tested LH stroke survivors, RH stroke survivors, and stroke-free control participants. For stroke participants, testing occurred around two as well as seven months after stroke, to capture neuroplasticity-induced changes between the sub-acute and chronic phase. The inclusion of RH stroke survivors further allowed us to investigate how hemispheric stroke location affects post-stroke lateralization of language-related functions. The EEG measurement was coupled with the CLaP in both sessions, covering aspects of language comprehension as well as production within each trial. Behavioral performance across sessions was characterized in terms of picture naming, as well as four neuropsychological tests tapping into non-linguistic and linguistic functioning (Boston Naming Test, Token Test, Amsterdam-Nijmegen Everyday Language Test, Corsi Block-Tapping Task). As mentioned above, scalp-level results can be majorly distorted by stroke lesions and should thus only be interpreted with caution (Piai et al., 2024; Piastra et al., 2022). To add to the interpretation of the EEG data, we performed individualized source-level analyses using the finite element method (FEM, Vorwerk et al., 2018) using session-specific individual structural MRI scans. This way we could control for lesion-induced distortions by precisely modelling the geometry and conductivity of the different tissues in individual brains, including the lesion.

## 2 Materials and Methods

This study was approved by the METC Regio Oost-Nederland (medical-ethics review committee, region east Netherlands) (NL72600.091.20, date: May 18, 2020), following the Code of Ethics of the World Medical Association (Declaration of Helsinki). Data collection took place at the Donders Centre for Cognitive Neuroimaging (DCCN, Radboud University) in Nijmegen, the Netherlands. Data and code will be made available via the Radboud Data Repository (https://doi.org/10.34973/1cyb-q622).

### 2.1 Participants and Recruitment

We recruited 14 control and 19 stroke participants. Inclusion criteria for stroke participants were either ischemic or hemorrhagic stroke of the middle cerebral artery in the left or right hemisphere, 18 to 78 years of age, right-handed native Dutch speaker, as well as sufficient communication abilities to give informed consent. Participants were included if they had a history of previous stroke if it did not cause any major or long-term impairments. For the control group, we recruited right-handed native Dutch speakers with no history of stroke or other brain disorders (self report), group-matched in sex and education to the stroke participants. Due to missing data of one control and one LH participant, the final sample consisted of 31 participants (C = 13, LH = 8, RH = 10).

We recruited stroke individuals via the neurology department of Radboud University Medical Center and the speech-and-language pathology department at the Canisius Wilhelmina Hospital in Nijmegen. All stroke participants were diagnosed by a neurologist based on clinical presentation and a neurological examination, supported by CT or MRI. The control group was recruited via participation in previous studies, social media advertisements, and social environment. Prior to participating, all participants received a study-specific information letter and gave written informed consent, and later received monetary compensation.

### 2.2 Materials

#### 2.2.1 Neuropsychological Tests

For a characterization of the sample, all participants underwent four neuropsychological assessments consisting of the Boston Naming Test (Kaplan et al., 1983; short version with 29 items based on Roomer et al., 2011), the Token Test (De Renzi & Vignolo, 1962; short version with 36 items), the Corsi Block-Tapping Task (Kessels et al., 2000), and both parallel versions of the Dutch and adapted (Ruiter et al., 2016, 2023) version of the Amsterdam-Nijmegen Everyday Language Test (ANELT, Blomert et al., 1995). The Boston Naming Test (BNT) evaluates confrontation naming abilities by means of black and white line drawings. The Token Test measures the overall severity of aphasia based on a number of tasks with tokens of different shapes, colors, and sizes. The Corsi Block Tapping task is a language-unrelated measure of visuospatial working memory where participants have to repeat an increasingly long sequence of blocks by finger tapping. The final score represents the highest number of blocks repeated correctly in one sequence multiplied with the number of correct trials (Kessels et al., 2000). The ANELT is a scenario test targeting verbal functional communication skills. To report a quantitative measure of verbal efficiency (i.e., average amount of essential information produced per time unit), we scored the ANELT based on the average number of essential content units (ECUs) produced per minute (Ruiter et al., 2011).

#### 2.2.2 Concise Language Paradigm

During the EEG recording, participants performed the Concise Language Paradigm (CLaP, Roos et al., 2024), consisting of auditory sentences followed by visual images to be named. Sentences were either contextually constrained, unconstrained or time-reversed, and images were photographs of objects (k = 126) or scrambled objects (k = 30). We employed auditory recordings from a female native Dutch speaker (Chupina et al., 2022), where sentence length varied from 4 to 8 words (M = 6) and audio duration was 1.79 to 3.59 seconds (M = 2.61, Mdn = 2.62). The cloze probabilities for the real-speech sentences (i.e., percentage of people completing the sentence with the intended target word) were 0-39% in unconstrained sentences and 60-100% in constrained sentences (*t*(77) = 51.236, *p* < .001) (Chupina et al., 2022; Hustá et al., 2021). The visual stimuli were derived from the BOSS database (Brodeur et al., 2010)(Brodeur et al., 2010) and the internet. To create scrambled stimuli, we distorted the normal photographs without majorly changing their basic visual properties (i.e., diffeomorphic transformation, see Stojanoski & Cusack, 2014).

### 2.3 Experimental Design

One EEG testing session consisted of 234 experimental trials presented in a uniquely pseudorandomized list per participant and session (using Mix, van Casteren & Davis, 2006). Comprehension conditions for the auditory sentences were constrained (k = 78), unconstrained (k = 78), and reversed speech (k = 78), and production conditions for picture naming (defined by the preceding sentence) were constrained (k = 78), unconstrained (k = 78), bare (k = 48), and scrambled (k = 30) pictures (Figure 1). Constrained and unconstrained pictures were always preceded by a matching sentence, while bare and scrambled pictures were always preceded by a reversed speech sentence. In sum, the CLaP yields contrasts of auditory sentence comprehension of real versus reversed speech during the sentence interval, contextually guided word retrieval during the pre-picture interval of constrained trials, and visually guided word retrieval during the picture interval of unconstrained, bare, and scrambled picture trials after picture onset.

**Figure 1.**
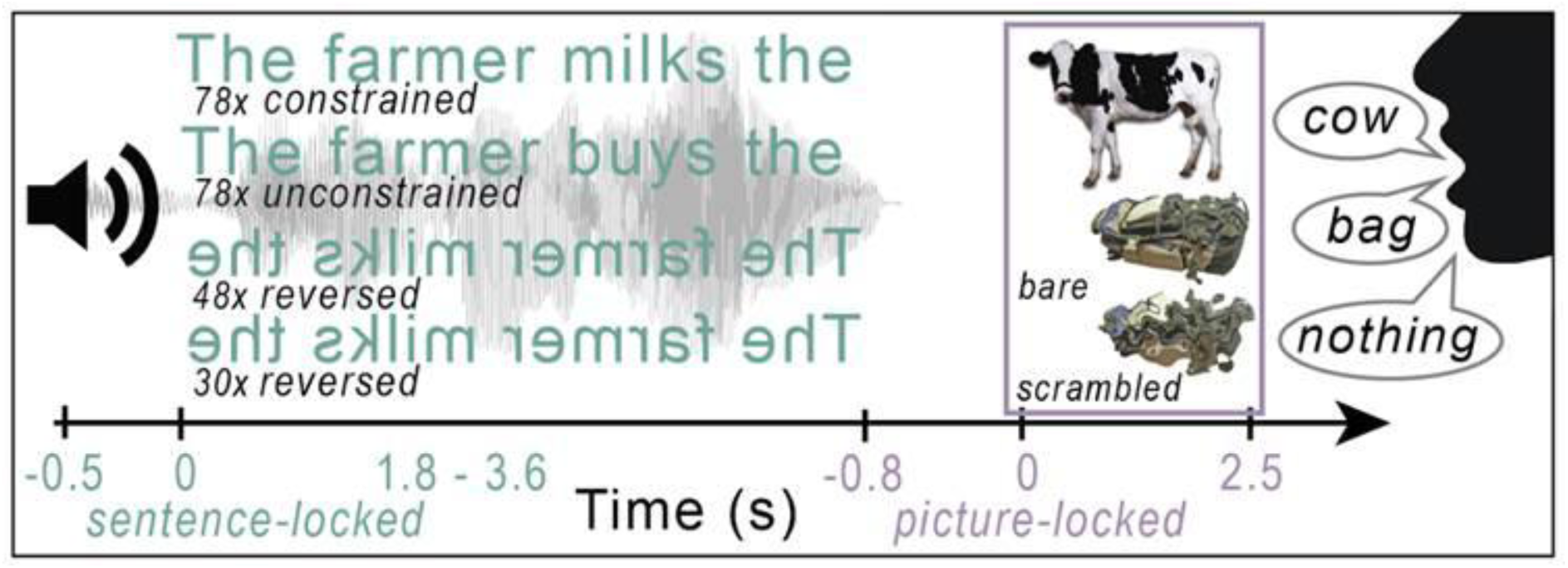
Schematic overview of example trials for all three sentence conditions (constrained, unconstrained, reversed) and both picture types (normal, scrambled), yielding constrained, unconstrained, bare, and scrambled picture naming conditions, as defined by the preceding sentence. Note the different (color-coded) time-lockings to sentence or picture onset. Sentence durations varied per trial. Reproduced with permission from the authors from https://osf.io/w5q7s/files/s68qr

### 2.4 Session Procedure

After clarifying remaining questions, participants signed the informed consent form and were screened for MRI compatibility. Sessions usually started with the MRI scan (ca. 20 min), followed by the four neuropsychological behavioral tests in a fixed order (BNT, Token Test, ANELT, Corsi). For the ANELT, we counterbalanced the starting version across participants and sessions, such that half of the participants started with version I and the other half with version II. The BNT and ANELT were audio recorded for later transcription and response scoring. We then started with the EEG preparation during which the participants were familiarized with the EEG task and the visual stimuli. We instructed them to listen to the auditory stimuli and name the images with the familiarized words. Scrambled pictures were named with the word “niks” (Dutch for *nothing*). We asked them to sit relaxed and try to keep blinks and movement to the designated blinking intervals and breaks. The experiment was presented on a 24-inch computer screen (Benq XL2420T, 1920×1080 resolution, 16:9 ratio) 60 to 80 cm from the participant using Presentation software (*Neurobehavioral Systems Inc*., Berkeley, CA, www.neurobs.com). Visual stimuli appeared centered on screen, 270 pixels in height, on a black background. We recorded picture naming responses from picture onset for 3.5 seconds using a microphone placed close to the mouth, for offline naming time and accuracy analysis.

Trials started with a centered fixation cross (1500 ms) followed by the auditory stimulus. After the pre-picture interval (800 ms), the visual stimulus was presented (2500 ms) followed by a blinking interval (2000 ms) with three asterisks (***). Every session started with practice trials of different conditions while the experimenter was still in the room to provide feedback. One EEG session consisted of nine blocks of 26 trials separated by breaks, and lasted about 40 minutes. With the end of the EEG recording, the testing session of about 3 hours was completed and we scheduled session 2 (equivalent to session 1) to take place five months later.

### 2.5 MRI Acquisition and Preprocessing

Participants had to be metal-free on their upper body to enter the MRI scanner, confirmed by a ferromagnetic detector. We placed vitamin-E pills on both ear lobes and the nasion as fiducial landmarks for co-registration of the MRI and the EEG data. We positioned participants on the scanner bed in a comfortable way with cushions and a tape over the forehead to minimize head movement. Scans were acquired with a 32-channel head coil, either on a 3-T Siemens Magnetom Prisma or a PrismaFit scanner. The MRI scanner choice depended on availability, but both sessions of one participant were always acquired on the same scanner. After a localizer and automatic alignment scan to adjust the field of view, we acquired high-resolution anatomical T1 images with a magnetization prepared rapid acquisition gradient echo (MPRAGE) (TR = 2300 ms, TE = 3.03 ms, 192 slices of 1 mm, voxel size = 1 mm isotropic, FOV = 256 mm, acquisition time = 5m21s), followed by an anatomical T2 fluid-attenuated inversion recovery (FLAIR) scan (TR = 5000 ms, TE = 397 ms, 192 slices of 1 mm, voxel size = 1 mm isotropic, FOV = 256 mm, acquisition time = 4m42s). DICOM images were converted to NIfTI files and bias-field corrected prior to registering all scans to the T1 image of each session for lesion delineation.

#### 2.5.1 Lesion Delineation and Statistics

For participants with stroke, we manually delineated the stroke lesions per session based on visual inspection of the structural MRI scans (T1 and T2), and sometimes the diffusion-weighted imaging. The lesion masks were drawn in itk-snap (www.itksnap.org) by visually detecting the lesion location and marking areas where voxel intensities abnormally differed from the healthy tissue surrounding the lesion in the different scans. These were then saved as binary masks in native space. Per participant, we extracted the voxel counts per segmented tissue type (see section 2.8.2, ‘Forward models’) and calculated the percentage of tissue damage based on grey and white matter combined. We examined the effects of group, session, and their interaction on lesion percentage using a linear-mixed effects regression model in R (R Core Team, 2017). To transform the lesion masks from native to MNI space, we used the SimNIBS tool *subject2mni*. Based on these normalized lesion masks and the Brainnetome atlas (Fan et al., 2016), we calculated the tissue damage per affected atlas parcel (in voxel counts and percentage), as well as the percentage of the lesion per damaged atlas parcel (i.e., how is the lesion distributed over the parcels).

#### 2.5.2 White Matter Hyperintensities

To assess white matter hyperintensities, we visually quantified the severity of periventricular and deep white matter hyperintensities (WMH) on all participants’ (including controls) T2 FLAIR MRI scans per session with a score from 0 to 3 (0 = no lesions, 1 = punctate lesions, 2 = beginning confluence, 3 = large confluent areas) based on the Fazekas scale (Fazekas et al., 1987). The Fazekas scale is the most widely used score to assess WMH indicative of small vessel disease and associated with cognitive decline (Tuladhar et al., 2016). Scoring was done by two raters independently per session. In cases where the two scores deviated, both raters came to a mutual agreement for the final score.

### 2.6 EEG Acquisition

We used ActiCap active electrodes (BrainProducts, Gilching, Germany) and EEG caps based on the equidistant M10 electrode layout (Easycap GmbH, Hersching, Germany), placed such that the central electrode (corresponding to Cz) was centered between nasion and inion and both ears. We recorded the locations of the electrodes by taking a 3D scan of the electrodes on the head, together with the fiducial points on the nasion and both ears marked by the vitamin-E pills from the preceding MRI scan. We then positioned participants in front of the screen in the electrically shielded EEG room and applied gel to each electrode. For the ground electrode (placed on the forehead) and the online reference electrode (equal to FCz) we aimed for an impedance below 5kOhm. For all other electrodes we aimed for an impedance below 25kOhm, which was not always possible due to time constraints or participant comfort. The EEG was recorded continuously with 64 active electrodes using BrainVision Recorder and an ActiChamp amplifier (500 Hz sampling rate, 0.1-200 Hz band-pass). For recording the horizontal and vertical electro-oculogram (EOG), we placed facial electrodes left of and underneath the left eye. Together with the respective counterpart electrode in the cap (right of the right eye, above the left eye) we computed two bipolar EOG signals during offline analysis. Two additional facial electrodes were placed above and below the right side of the lips to detect speech-related activity to facilitate artifact identification. We recorded stimulus markers synchronous with the EEG at the onset of each sentence and picture.

### 2.7 Picture Naming Times and Accuracy

We scored the accuracy of picture naming responses for the CLaP either online or post-hoc based on the trial recordings and manually measured response times (RT) from picture onset using PRAAT (Boersma & Weenink, 2017), naive to condition. Response onset was always marked at the target word (if included) or a close synonym. Hesitations or self corrections are thus reflected in longer response latencies. If the response did not include the target word or a synonym, the respective trial was marked as incorrect and excluded from RT analysis.

We statistically evaluated RTs and error rates in R (R. Core Team, 2017) using generalized linear mixed-effects regression (package lme4, Bates et al., 2015) with fixed effects of condition (four levels), group (three levels), and session (two levels). The linear model for RTs contained all two-way and three-way interactions as well as by-participant and by-item random intercepts. For errors, a logistic regression was used with each of the three fixed effects and an interaction between group and session, with by-participant random intercepts (more complex models failed to converge). All variables underwent deviation coding. The estimates for the relevant level contrasts were subsequently obtained from the RT and error rate models using emmeans (Lenth, 2023).

### 2.8 EEG Preprocessing

Unless otherwise noted, we performed all EEG preprocessing and analysis steps in MATLAB (version R2024b) using the FieldTrip toolbox (version 20250526, Oostenveld et al., 2011). As for the RT analysis, we only included correct trials for the EEG analysis, with the exception of hesitations and self-corrections, which were marked as correct for RT and error analyses, but excluded from EEG analyses. We cleaned the data on the basis of whole trials (from 800 ms prior to sentence onset until 400 ms after picture onset). All trials were low-pass filtered (50 Hz, twopass Butterworth IIR filter with order 6) and baseline corrected using the 800 ms interval before sentence onset. Based on visual inspection, we first rejected channels which remained noisy for the majority of the recording and then performed Independent Component Analysis (ICA) to reject eye-movement and blinking components from the data (Jung et al., 2000). To improve the detection of these components by the ICA, we rejected extremely noisy trials from the data on which the ICA was performed, but rejected the components from the data including all trials.

After ICA, we further rejected noisy channel segments per trial based on signal variability. We calculated each channel’s standard deviation (SD) per trial and defined a global, session-specific rejection threshold as the median SD across all channels and trials plus two median absolute deviations (MAD) of these SDs. Channels whose SD exceeded this threshold in a given trial were removed from that trial. Trials in which this led to a rejection of more than 50% of the channels were fully rejected. All other missing channel segments were interpolated based on the average of the neighboring channels. Finally, before interpolating the fully rejected channels, we split the data for scalp and source analysis. Source analysis data was only re-referenced to the common average of all channels, while scalp analysis data was first interpolated to add the rejected channels and then re-referenced to the common average.

### 2.9 EEG Analysis

In the following, we report our methodological approach separately for scalp and source level. For both levels, these are divided into three contrasts throughout the course of each trial (Figure 1), consisting of auditory sentence comprehension (sentence interval), contextually guided word retrieval (pre-picture interval), and visually guided word retrieval (picture interval). Overall, the present scalp-level analyses were based on previous findings for the CLaP from a group of young, stroke-free control participants (Roos et al., 2024). Source-level analyses were used in addition, to add to the interpretation of the results of the stroke groups (Piai et al., 2024). These were mainly informed by the scalp-level (and source-level) results of the control group, reported in detail below.

#### 2.9.1 Scalp-level Analysis

We expected the scalp-level results (of the control group) to be in line with previous findings for the CLaP (Roos et al., 2024). We therefore analyzed the data accordingly by comparing event-related potentials (ERPs) or time-frequency-representations (TFRs) between conditions per contrast, separately per session and group (C, LH, RH).

The general approach for all contrasts consisted of equating the number of trials per comparison (max. 78 for constrained, unconstrained, or reversed, max. 30 for bare and scrambled). To evaluate the consistency of these effects across sessions, we compared the differences in amplitude (ERPs) or normalized differences in power (TFRs) of each contrast across sessions. The differences between conditions and sessions were all statistically evaluated by a cluster-based permutation test including all channels (Maris & Oostenveld, 2007). Dependent-samples *t*-tests were computed across participants, and clusters of neighboring electrodes (minimum two) surviving a threshold of *p* < .05 were identified. Cluster-level statistics were calculated as the sum of *t*-values within each cluster and evaluated using a two-tailed permutation test with *α* = .05. Analysis steps that were contrast specific are reported below.

##### Sentence Comprehension

We previously found that both constrained and unconstrained sentence ERPs significantly differed from those of reversed speech sentences in the first 500 ms after sentence onset (Roos et al., 2024). Here, we further simplified this contrast by averaging the ERPs of constrained and unconstrained sentences to gain a measure of real speech, and compared it to the ERPs of reversed speech sentences from −800 to 1800 ms relative to sentence onset (length of the shortest included sentences).

Per participant, sentence ERPs were low-pass filtered at 40 Hz, baseline-corrected with the 800 ms prior to sentence onset. Session-consistency was evaluated by means of the difference wave for real minus reversed speech across sessions per group.

##### Pre-Picture: Contextually guided word retrieval

The context effect, that is contextually (constrained) versus visually (unconstrained) guided word retrieval, has been repeatedly shown to lead to power differences in the alpha-beta frequency range prior to picture onset (Klaus et al., 2020; Piai et al., 2015, 2017, 2018; Roos et al., 2024; Roos & Piai, 2020). We thus analyzed the difference between constrained and unconstrained trials during the 800 ms pre-picture interval by means of TFRs per group and session.

Per participant, TFRs were computed from 3-40 Hz with a sliding time window of 3 cycles advancing in steps of 50 ms and 1 Hz. Each time window was multiplied with a Hanning taper to control for spectral leakage. Across-session consistency was tested by statistically comparing the power differences between constrained and unconstrained sentences across sessions per group.

##### Picture: Visually guided word retrieval

In a previous study employing the CLaP (Roos et al., 2024), we investigated ERPs for object recognition and naming across all four picture conditions (constrained, unconstrained, bare, scrambled) and visual evoked potential components (P1, N, P2). In the present study, the theoretically motivated contrasts consisted of P2 amplitudes (see for discussion Piai & Borges, 2023; Roos et al., 2024) for constrained versus unconstrained pictures (tapping into spreading activation during word planning) as well as bare and scrambled pictures (tapping into word planning while controlling for general visual and motor effects). Across-session consistency of these comparisons was statistically evaluated by the difference wave of constrained minus unconstrained and bare minus scrambled on group-level per session.

Picture ERPs were low-pass filtered at 40 Hz. Note that picture-locked ERPs could not be baseline-corrected using a time window just before picture presentation, as they were preceded by different sentence conditions leaving no common and unbiased baseline window for such a correction. Given the systematic differences preceding picture presentation, our analyses included the entire interval of −1000 to 400 ms relative to picture onset.

#### 2.9.2 Source Level Analysis Forward models

For the reconstruction of the underlying sources of the activity measured on the scalp by means of inverse modelling, we first constructed individual forward models per participant and session, consisting of the volume conductor model, the source model, and the electrode configuration.

The volume conductor model is a volumetric, three-dimensional geometrical representation of the head anatomy, including the electrical conductivity of its different tissue types. To compute this, we first constructed high-resolution tetrahedral tissue meshes based on the T1 and T2-weighted structural MRI scans using SimNIBS (version 4.0.0). The SimNIBS *charm* tool (Puonti et al., 2020) segmented the scans into the following tissue types: white matter, grey matter, csf, scalp, eye balls, compact bone, spongy bone, blood, muscle. Where necessary, we manually corrected these tissue segmentations and added the binary lesion masks for the participants with stroke as an additional tissue (function: add_tissues_to_upsampled). We then re-meshed the segmentation to obtain one volumetric mesh per tissue type with smooth boundaries. We then added the electrical conductivity for each tissue type in Siemens per meter (white matter = 0.22, grey matter = 0.47, csf = 1.71, scalp = 0.41, eye balls = 0.5, compact bone = 0.005, spongy bone = 0.048, blood = 0.57, muscle = 0.32, lesion = 0.75, based on McCann et al., 2019) and used the finite element method (FEM) based on the SimBio software as implemented in FieldTrip (Vorwerk et al., 2018) to compute the volume conductor model.

The source model consists of the locations of the equivalent current dipoles in the brain that could have generated the electrical activity measured on the scalp. To build the source model, we first placed dipoles at each centroid of the high-resolution tetrahedral mesh for the grey matter, resulting in a variable number of dipoles per participant. We also constructed an MNI-template source model with a predetermined resolution of 3 mm, resulting in a grid of 36,271 dipoles in the grey matter. We warped this grid to the individual high-resolution source model and moved every dipole to the closest centroid of the individual grey matter mesh, resulting in approximately 3 mm resolution source models with the same number of dipoles across participants.

The electrode configuration was recorded using a 3D structure sensor (Occipital Inc., Boulder, CO, www.structure.io) mounted on an Apple iPad (Apple, Cupertino, CA) prior to the EEG measurement (Homölle & Oostenveld, 2019). We coregistered these 3D structures to the MRI scans by means of the three fiducial landmarks recorded in both scans (nasion, left and right ear lobe), and then manually located the position of each electrode in 3D space relative to the fiducials. The co-registration of the electrode structure to the MRI scan was confirmed by plotting both structures in one figure and adjusting their alignment by rotating and/or translating the electrode structure to obtain the maximum fit. For two participants, we could not acquire a 3D structure of the electrode positions at one of the sessions due to technical equipment failures, in which cases we used the 3D electrode configurations of the respective other session of the same participant. As the EEG cap was always placed in the same manner (see section 2.6), we do not expect major deviations for the co-registration between the two sessions.

With the volume conductor model, the source model, and the coregistered electrode configuration, we then computed the lead field (forward model).

#### 2.9.3 Source Reconstruction

The source-level analyses were based on the scalp-level results of the control group, as well as previous source-level findings (Piai et al., 2015, 2017, 2018; Roos & Piai, 2020). We applied different source reconstruction techniques per contrast, as detailed below, but will first present the general framework for analyses of all contrasts.

##### Determination of ROIs

As the control group showed no changes between sessions on scalp level, we averaged the source results of both sessions per contrast. We then statistically evaluated the difference between conditions at each grid point (based on template positions) across controls using dependent-samples *t*-tests to define regions of interest (ROIs). These ROIs were determined by Brainnetome atlas parcels (Fan et al., 2016), such that any parcel containing grid points below a contrast-specific alpha-threshold entered the ROI map. Per contrast, this threshold was determined by statistically evaluating the differences between conditions using a parametric voxel-wise dependent samples t-test. To capture laterality changes to homologous areas in the stroke participants, as reported in previous literature (Crinion & Price, 2005; Saur et al., 2006; Wilson & Schneck, 2021), the ROIs were mirrored at the midline. In other words, the ROIs included all parcels exhibiting differences between conditions below the specified threshold *bilaterally*, regardless of the hemisphere in which the sources were reconstructed in the control group.

##### Sentence Comprehension

Following the scalp-level analysis, we source reconstructed the differences between real and reversed speech between 300 and 450 ms relative to sentence onset per group. As beamforming techniques assume temporally uncorrelated sources (Westner et al., 2022), a minimum norm estimation (MNE) solution was preferred for the source reconstruction of the auditory ERP peak. Bilateral auditory sources are usually highly correlated in time, in which case beamforming tends to perform poorly (Diwakar et al., 2011; Murzin et al., 2013). We used the MNE implementation of FieldTrip (Dale et al., 2000; pre-whitening, lambda regularization = 3, scaled source covariance matrix, fixed source orientation) and averaged the resulting time-resolved source power values over time per participant. The statistical comparison in the control group was evaluated at *p* < .001 (uncorrected) and all parcels containing grid points satisfying this threshold entered the ROI map for sentence comprehension.

##### Pre-Picture: Contextually guided word retrieval

Based on previous studies, we expected the context effect to show in the alpha-beta frequency range during the pre-picture interval (Roos & Piai, 2020) and hence used a Dynamic Imaging of Coherent Sources (DICS) beamformer (Gross et al., 2001; unit-noise-gain normalization, lambda regularization = 5, fixed source orientation). First, we computed the complex Fourier spectra during the 800 ms preceding picture onset using multitaper time-frequency transformation at 12 Hz with spectral smoothing of plus and minus 4 Hz. The common spatial filters were computed on the complex Fourier spectra of both constrained and unconstrained sentences together, and then applied to the Fourier spectra of constrained and unconstrained sentences separately. For controls, We then statistically compared constrained and unconstrained pre-picture intervals and all parcels containing grid points satisfying a threshold of *p* < .001 (uncorrected) entered the ROI map for contextually guided word retrieval.

##### Picture: Visually guided word retrieval (P2)

For the picture interval we investigated the differences between constrained and unconstrained as well as bare and scrambled pictures during the P2 ERP component from 250 to 350 ms after picture onset, informed by the control group scalp-level results. As we did not expect sources to be correlated in time, we used a Linearly Constrained Minimum Variance (LCMV) beamformer (van Drongelen et al., 1996; Van Veen et al., 1997; unit-noise-gain normalization, lambda regularization = 5, fixed source orientation) with common spatial filters including all conditions from −150 to 350 ms relative to picture onset. For both contrasts (constrained minus unconstrained, bare minus scrambled), the statistical comparison in the control group was evaluated at *p* < .01 (uncorrected), and all parcels containing grid points satisfying this threshold entered the respective ROI map. This change in threshold relative to the other contrasts was necessary as the more stringent threshold did not yield suprathreshold grid points. For both P2 contrasts, however, this more lenient threshold yielded results in line with previous literature (Levelt et al., 1998; Liljeström et al., 2009; Sörös et al., 2003). Given that this is not the inferential statistical step of interest, this change in threshold affects the amount of measurement error introduced, but it does not affect the strictness of the inferential step in the analysis of the stroke groups.

##### Changes in hemispheric dominance

For the stroke participants, we first summarized the differences between conditions over trials on the individual level using independent-samples *t*-tests. We then focused on evaluating changes in laterality or hemispheric dominance between the two time points. Per participant and contrast (sentence, pre-picture, picture), we first averaged the resulting grid point *t*-values within each parcel belonging to the ROI map and then averaged these per hemisphere, to gain an ROI-based averaged t-value per hemisphere. This enabled us to quantify changes in hemispheric dominance for each stroke participant over time.

The individual laterality index (LI) was calculated per session as the difference between the absolute left and right hemisphere t-values divided by their sum, yielding values between −1 and +1, where positive values indicate a LH-dominance and negative values a RH-dominance. First, using a linear model, we verified that lesion size in session 1 did not systematically co-vary with hemispheric dominance in session 1, in interaction with group. Finally, we examined whether hemispheric dominance changed between sessions, using a generalized linear mixed-effects logistic regression. The binary outcome variable indicated whether hemispheric dominance changed between sessions (yes/no). Fixed effects included lesioned hemisphere (group: LH, RH, deviation coding), hemispheric dominance in session 1 (deviation coding), their interaction, and contrast type (simple coding). We included random intercepts for participants to account for repeated measures within individuals. Note that we did not explicitly test whether changes in hemispheric dominance differed per contrast type, as our stroke participant samples were relatively small.

## 3 Results

### 3.1 Sample Characteristics

Baseline characteristics of the study sample per group are listed in Table 1, and the results of the neuropsychological tests in Table 2. As the severity of WMH per participant did not change between sessions (regardless of age, group, or stroke type), we report one score per participant. For stroke participants, this score is an indication of premorbid brain state in terms of periventricular and deep white matter lesions.

**Table 1.**
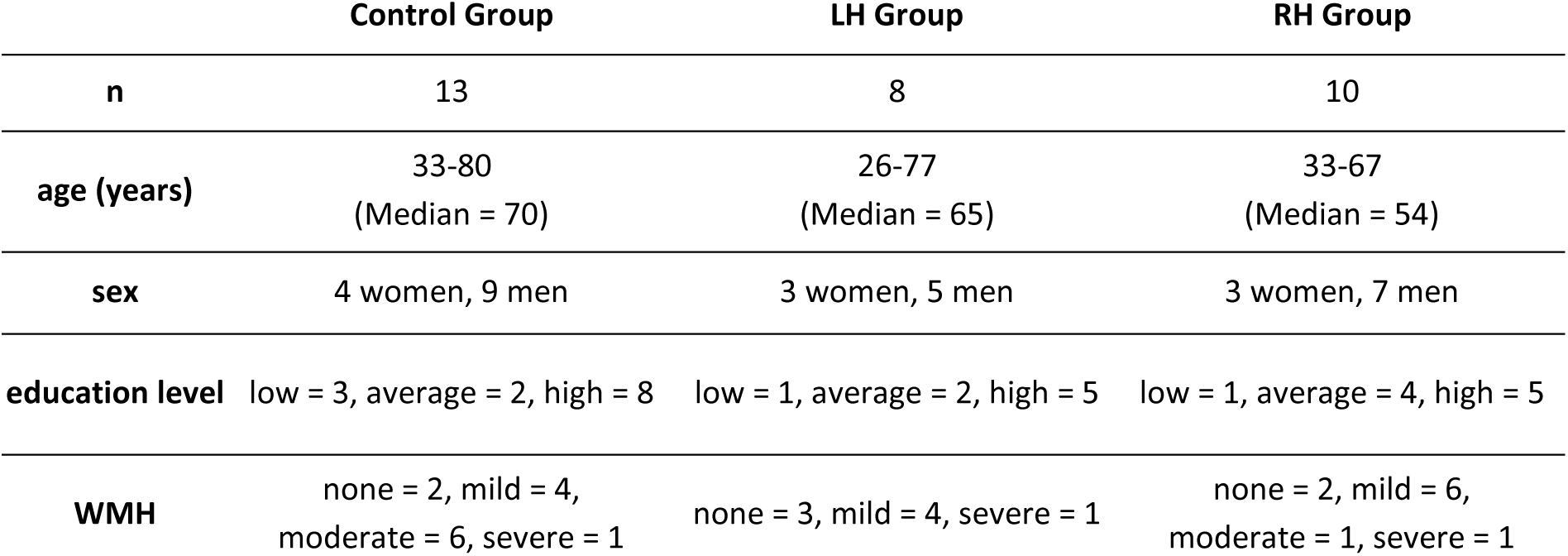
Characterization of the study sample in total as well as per group. Level of education is based on Verhage (1983). Education was quantified on a scale from 1 to 7 (Verhage, F., 1983) grouped into low (1-4), average (5) and high (6-7), and white matter hyperintensities (WMH) on a scale from 0 to 3 (Fazekas et al., 1987) grouped into none (0), mild (1), moderate (2), and severe (3), reflecting the severity of periventricular and deep WMH per participant based on the MRI T2 FLAIR scans.

|  | Control Group | LH Group | RH Group |
| --- | --- | --- | --- |
| **n** | 13 | 8 | 10 |
| **age (years)** | 33-80<br>(Median = 70) | 26-77<br>(Median = 65) | 33-67<br>(Median = 54) |
| **sex** | 4 women, 9 men | 3 women, 5 men | 3 women, 7 men |
| **education level** | low = 3, average = 2, high = 8 | low = 1, average = 2, high = 5 | low = 1, average = 4, high = 5 |
| **WMH** | none = 2, mild = 4,<br>moderate = 6, severe = 1 | none = 3, mild = 4, severe = 1 | none = 2, mild = 6,<br>moderate = 1, severe = 1 |

**Table 2.** Mean test scores and standard deviation in parentheses per neuropsychological test for all groups and both sessions.

|  | Control Group |  |  |  | LH Group |  |  |  | RH Group |  |  |  |
| --- | --- | --- | --- | --- | --- | --- | --- | --- | --- | --- | --- | --- |
|  | BNT | Token | Corsi | ANELT | BNT | Token | Corsi | ANELT | BNT | Token | Corsi | ANELT |
| **session 1** | 162.8<br>(7.3) | 33.8<br>(2.5) | 40.8<br>(9) | 31.8<br>(14.1) | 153.9<br>(9.8) | 31.8<br>(2.7) | 40.9<br>(7.4) | 20.2<br>(6.4) | 160.8<br>(11.4) | 34.7<br>(2.1) | 50.8<br>(20.4) | 23.7<br>(10) |
| **session 2** | 164<br>(6.8) | 33.9<br>(2) | 44.9<br>(14.2) | 32.1<br>(13.3) | 159<br>(13.4) | 32.2<br>(3.3) | 52.6<br>(11.4) | 25.2<br>(10.8) | 161.3<br>(11) | 35<br>(1.2) | 54.6<br>(24.1) | 33.2<br>(14.5) |

Note: Reported scores are raw test scores, not corrected for age or education. ANELT scores reflect verbal efficiency and BNT scores are corrected to the long version of the test (impairment cut-off: 149.6). Aphasia cut-off Token Test: 29.

Descriptively, the LH group obtained the lowest scores in all neuropsychological tests in session 1, followed by considerable improvement over time, approaching the scores of the other two groups in session 2 (Table 2). While participants in the RH group did not suffer from language impairments, the severity of language impairments in the LH group was mild at most.

#### Lesion Characteristics

Figure 2 shows the extent of LH and RH lesions per group and session. We evaluated the effects of group, session, and their interaction on lesion size in percentage (i.e., individual lesion size relative to individual brain size based on GM and WM voxels) using a linear-mixed effects regression model. There was no significant effect of Group (Estimate = −0.096, *SE* = 0.410, *t*(20.62) = −0.24, *p* = .817), no significant effect of Session (Estimate = −0.134, *SE* = 0.276, *t*(14) = −0.49, *p* = .634), and no significant Group by Session interaction (Estimate = 0.245, *SE* = 0.367, *t*(14) = 0.67, *p* = .515). This shows that lesion percentage did not significantly differ between groups or sessions.

**Figure 2.**
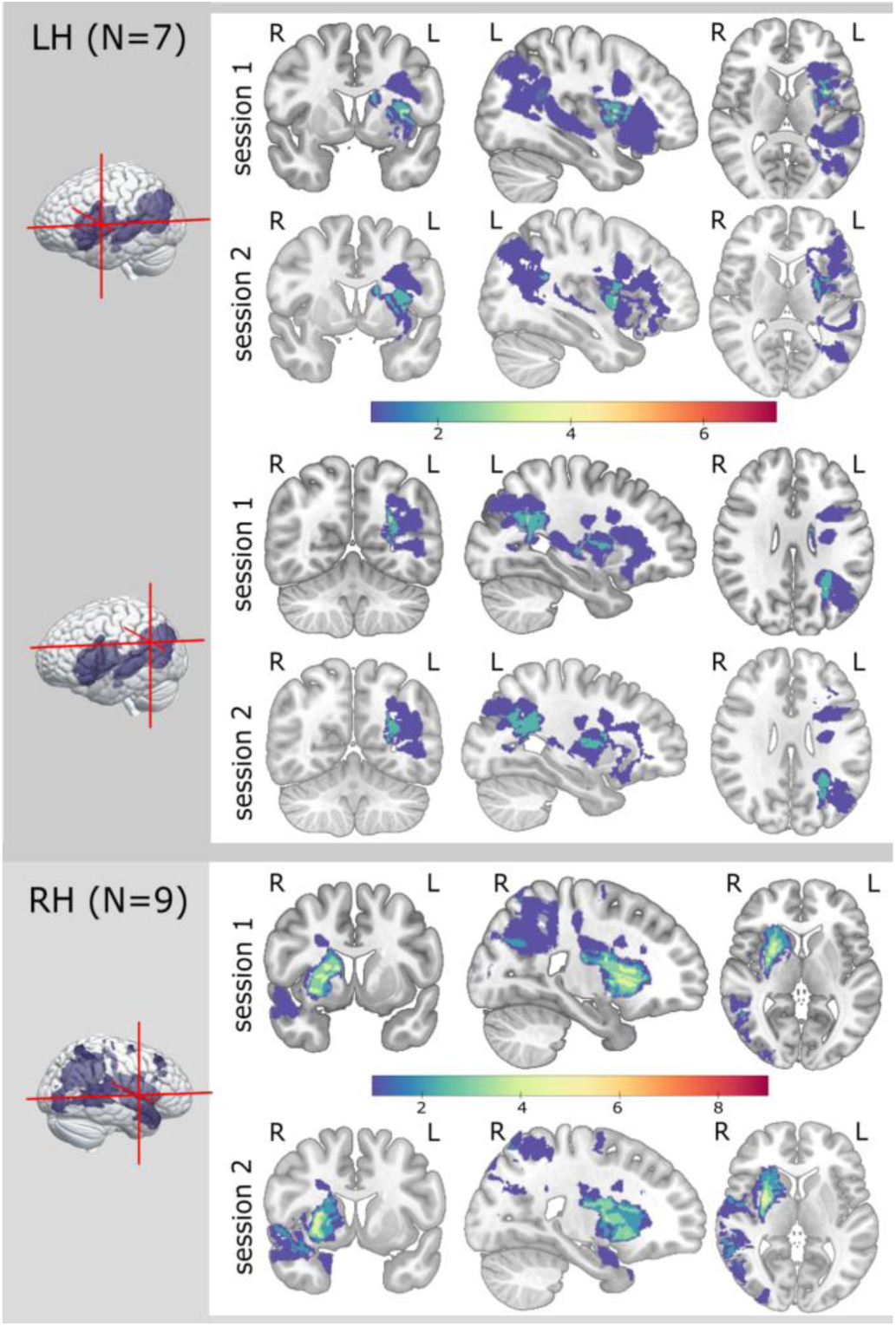
Lesion overlap maps per group and session based on manually delineated native-space lesion masks normalized to MNI space. Plotted using MRIcroGL (v1.2.20220720, Rorden, 2025). Note that the lesion masks per group (N) equal total group-size (LH = 8, RH = 10) minus 1, as both groups included one participant with imaging-confirmed stroke, but with no visually detectable lesion anymore two and seven months after stroke.

### 3.2 Picture Naming Times and Accuracy

The group-level error rates per session and condition are listed in Supplementary Table 1. Inferential statistics indicated that conditions and groups systematically differed from their overall mean (ps < .02). Group and session interacted (p < .001). The error rate was highest for the LH group in session 1 (5.9%), followed by improvements in session 2 (2.9%), as confirmed by the inferential statistics (estimate = .76, SE =.17, z = 4.47, p < .001). The RH group, by contrast, had a lower error rate in session 1 (2%) and in session 2 (2.4%), with no statistically significant differences between sessions (estimate = -.20, SE =.20, z = 1.01, p = .311). The same pattern was found for the control group, with no statistically significant differences between session 1 (1.3%) and session 2 (1.3%; estimate = -.05, SE = .22, z = -.23, p = .817). Interestingly, the more detailed classification of non-target responses in the LH group illustrates a shift from incorrect responses (3.3% in session 1, 1% in session 2) to correct synonyms or self-corrections (6.9% in session 1, 8.1% in session 2). Note that the latter non-target responses were marked as correct for the RT and error analysis (but not the EEG analysis).

Mean group-level picture naming RTs per condition are displayed per session and condition in Figure 3, and listed in Supplementary Table 2. Descriptively, all three groups showed the same pattern of picture naming times as previously established, such that constrained pictures were named the fastest, followed by scrambled pictures, and unconstrained and bare pictures (Roos et al., 2024). Following inferential statistics, conditions, groups, and sessions differed systematically from their overall mean (*p*s < .009). Group and session interacted (*p*s < .001), as well as group and condition (*p*s < .043). Regarding the simple effects within group, all pairwise comparisons of condition were statistically significant, except for the bare versus unconstrained comparison in the RH group (Supplementary Table 3). Controls did not differ in RTs between the two sessions (estimate = 4.43, SE = 7.44, z = 0.60, p = .551), while both stroke groups became faster in session 2 relative to session 1, with the LH group showing the largest improvement (LH: estimate = 84.84, SE = 9.72, z = 8.731, p <.001; RH: estimate = 17.42, SE = 8.51, z = 2.05, p = .041; all p-values adjusted with Tukey method).

**Figure 3.**
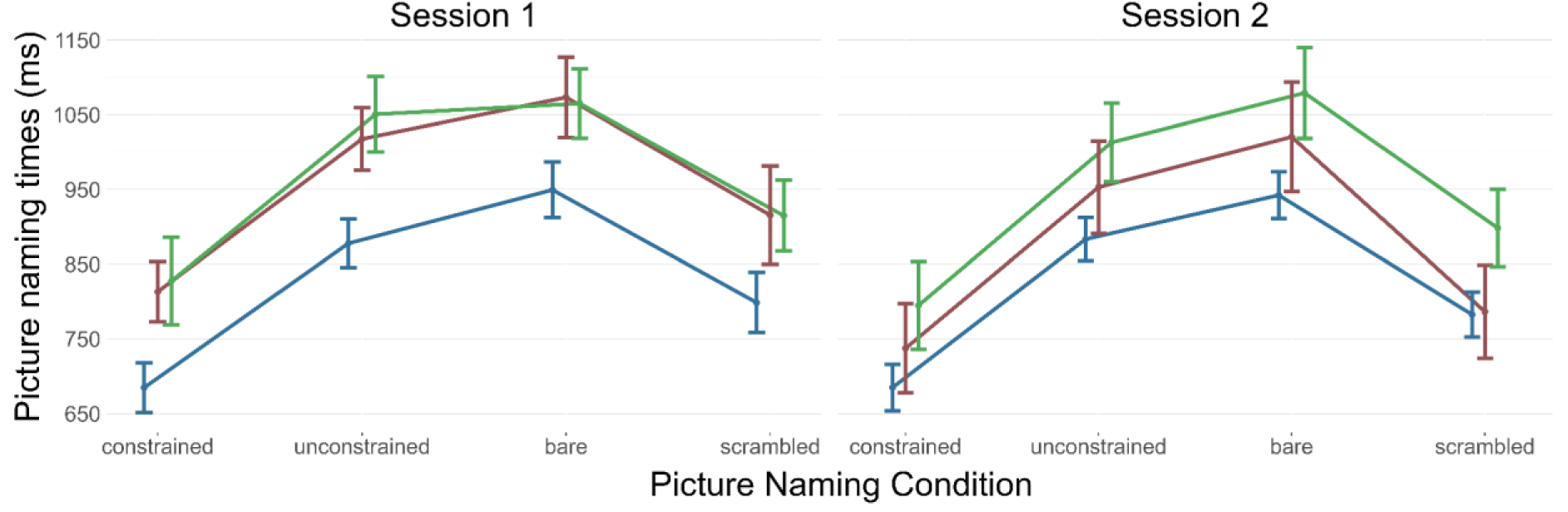
Mean picture naming times (ms) per picture condition for session 1 (left) and 2 (right) for all groups (C = blue, LH = red, RH = green). Error bars show standard error of the mean.

In sum, across naming RTs and error rates, the LH group showed improvement between session 1 and session 2, while the improvement in the RH group was modest and only present for the RTs. The stroke-free control group, as expected, remained stable.

### 3.3 EEG Results

Below, we report the scalp and source-level EEG results, divided into sentence comprehension, contextually guided picture naming (pre-picture interval) and visually guided picture naming (picture interval). Per contrast, we first report the scalp-level results, which we expected to be in line with previous findings from the CLaP for the same contrasts (Roos et al., 2024). Then we turn to the source-level findings, where we report the session-averaged results of the control group in terms of ROI maps based on grid-point alpha-thresholds used to assess changes in hemispheric dominance in the stroke participants. Finally, we address the changes in hemispheric dominance over time for the stroke participants.

#### 3.3.1 Sentence Comprehension

During the sentence interval, we detected stronger evoked peak amplitudes for real compared to reversed speech ERPs (Figure 4), in line with Roos et al. (2024). These amplitude differences occurred between 300 and 450 ms after sentence onset and were statistically significant across groups and sessions (ps < .008, except for LH session 2, where p < .02). For the control group in session 2, we observed an additional cluster of amplitude differences (p = .0017) during the course of real versus reversed speech sentences (cluster associated with time points 806 to 1120 ms).

**Figure 4.**
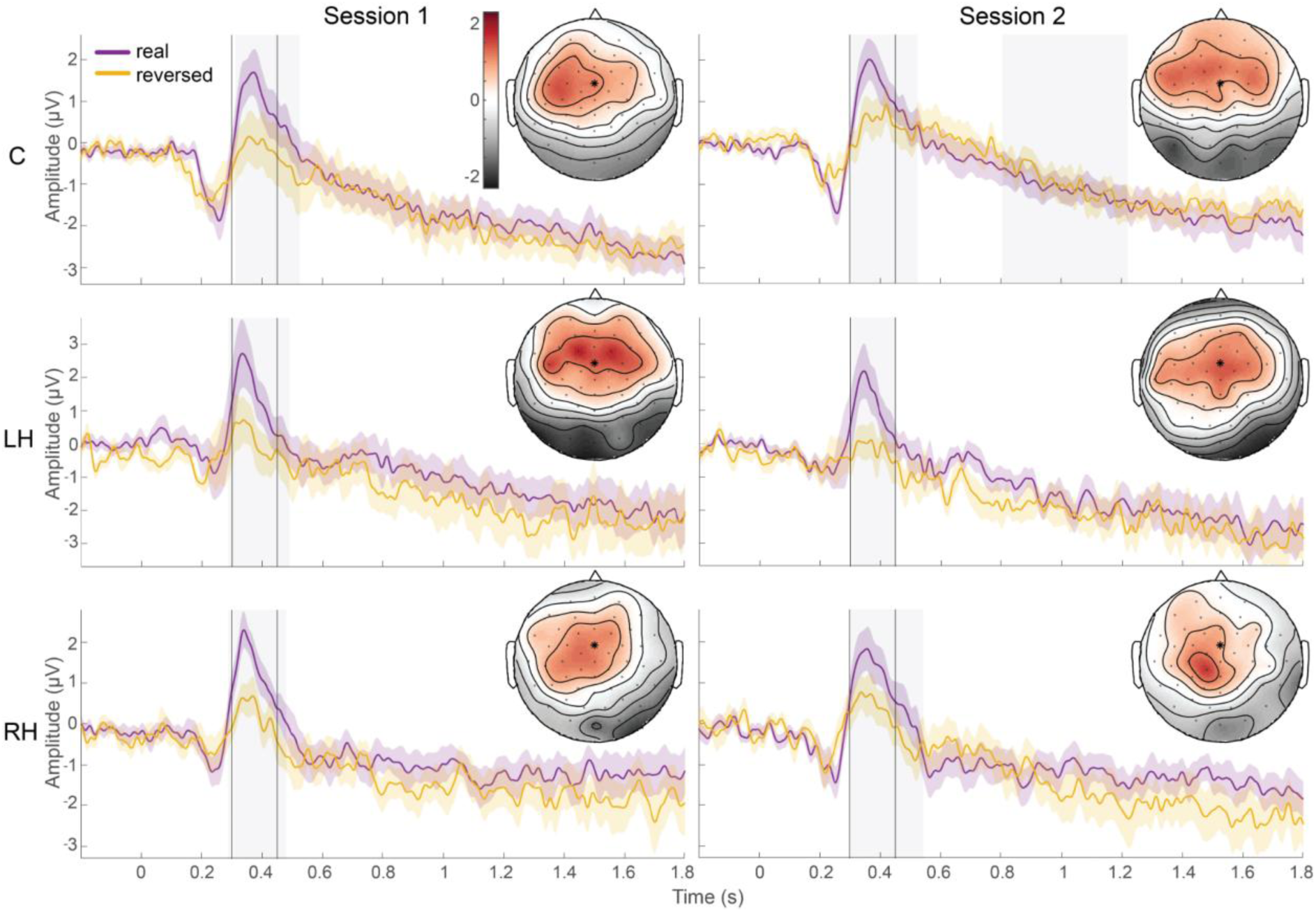
Scalp ERPs from −200 to 1800 ms relative to sentence onset for real (purple) and reversed speech (orange) for one central electrode per session (columns, session 1, session 2) and group (rows, C: control, LH: left hemisphere, RH: right hemisphere). The location of the electrode is marked in the topographic plots with an asterisk (comparable to FCz). Color-shaded areas around the waveforms represent standard error of the mean. Grey shaded areas mark time intervals for real minus reversed speech per plot underlying the most prominent clusters within each comparison. Corresponding topographies show the difference between real and reversed speech from 300 to 450 ms after sentence onset, marked by two vertical lines in the ERP plot. Time 0 s represents sentence onset in all plots. Note that all topographies are scaled to the same range, but waveforms are scaled per group.

The across-session comparison per group based on the difference waves of real minus reversed speech ERPs yielded no significant differences, indicating no systematic differences in amplitudes between session 1 and 2.

Sources of the auditory ERP responses between 300 and 450 ms after sentence onset are shown in Figure 5A. The session-averaged differences between real and reversed speech sentences in the control group below *p* < .001 were reconstructed almost exclusively in LH areas, with the exception of the primary auditory cortex in the right superior temporal lobe. LH areas were distributed over the left temporo-parietal and frontal cortex, including posterior middle and inferior frontal gyrus, temporal lobe and fusiform gyrus, inferior parietal lobule, postcentral gyrus, insula, hippocampus, thalamus, as well as the basal ganglia (pallidum, caudate, putamen).

**Figure 5.**
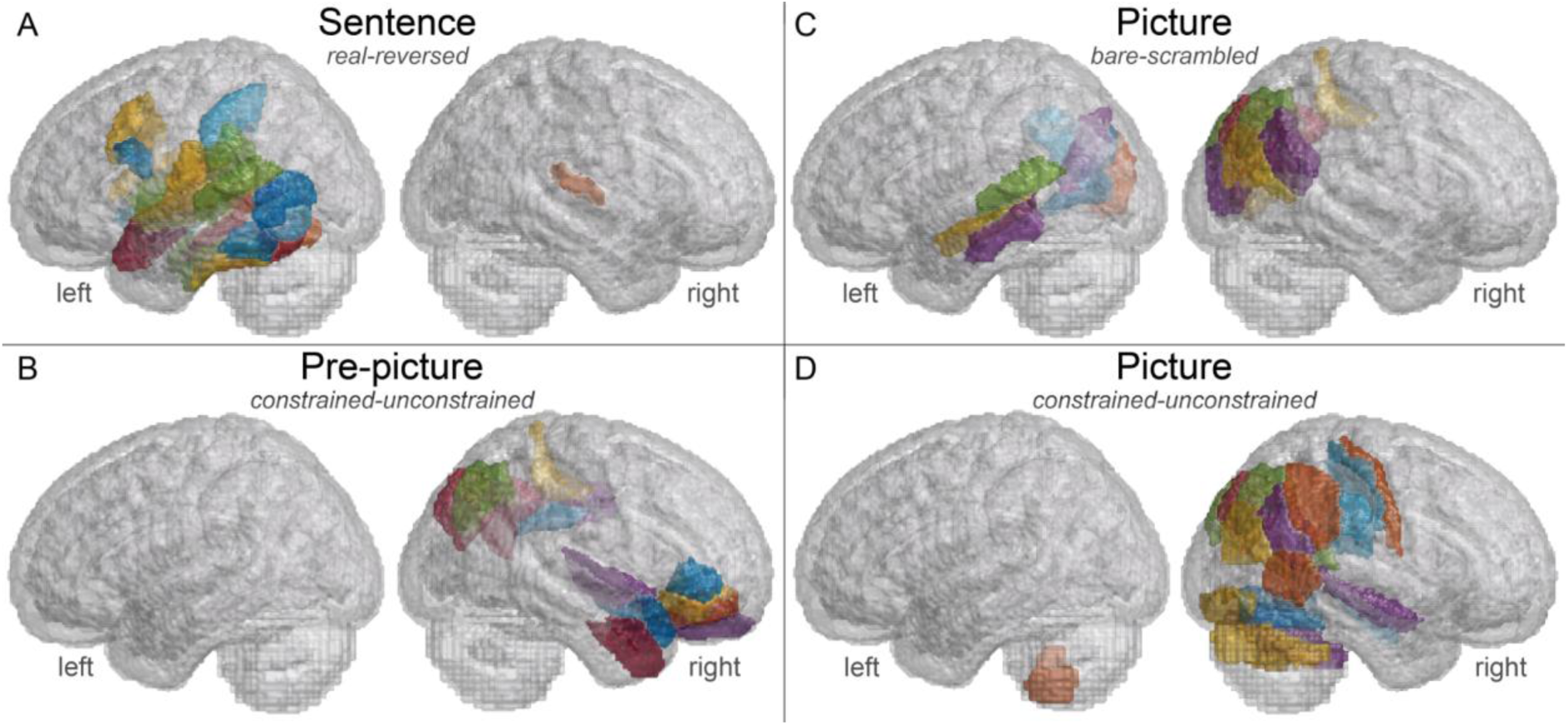
ROI maps per contrast based on session-averaged control group results showing Brainnetome parcels containing grid points below contrast-specific alpha-level thresholds. Sentence: MNE for real minus reversed speech from 300-450 ms after sentence onset, p < .001. Pre-picture: DICS for constrained versus unconstrained pre-picture intervals from −800 to 0 ms relative to picture onset centered at 12 Hz (8-16 Hz), p < .001. Picture: LCMV during P2 peak from 250-350 ms after picture onset for bare-scrambled pictures (top) and constrained minus unconstrained pictures (bottom), p < .01. All contrasts are plotted with the same parcel-color-scheme on a semi-transparent brain mesh. Same parcels thus appear in the same color across contrasts and deep grey matter structures (i.e., more medial) are less colorful. Note that cerebellar regions are based on the AAL atlas (Tzourio-Mazoyer et al., 2002), as the Brainnetome atlas (Fan et al., 2016) does not include the cerebellum.

#### 3.3.2 Pre-Picture: Contextually guided word retrieval

The context effect, tapping into contextually guided word retrieval, was associated with power decreases between 3 and 35 Hz during the pre-picture interval across groups and sessions (Control: session 1 p = .015, session 2 p = .043; LH: session 1 p = .0234, session 2 p = 0; RH: session 1 p = .08, session 2 p = .041), revealing the expected effect for contextually guided naming (Roos et al., 2024; Roos & Piai, 2020), depicted in Figure 6.

**Figure 6.**
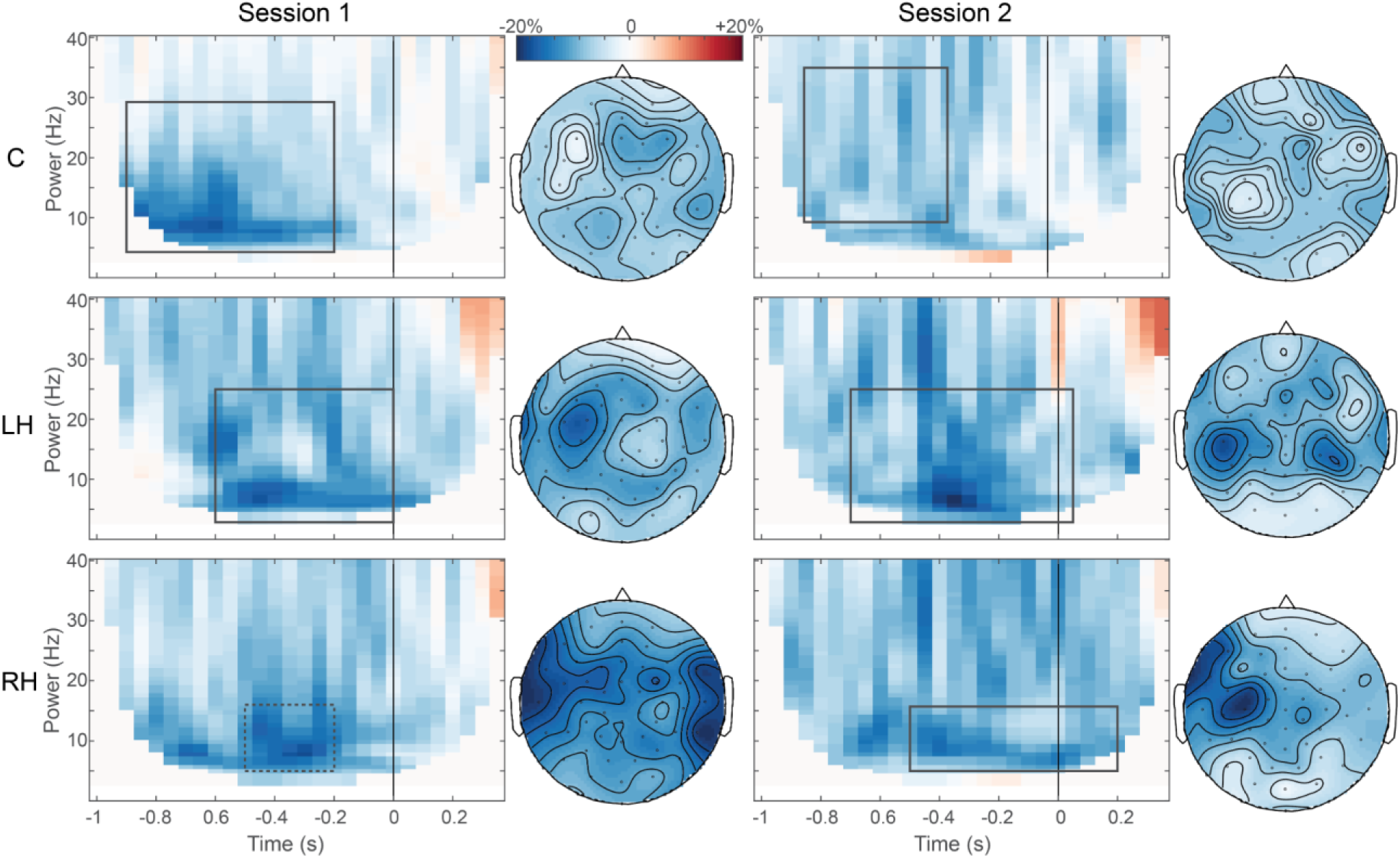
Scalp TFRs showing relative power changes from −1000 to 400 ms relative to picture onset for the context effect between constrained and unconstrained trials averaged over all electrodes per session (columns) and group (rows, C: control, LH: left hemisphere, RH: right hemisphere). Topographies correspond to most prominent time and frequency intervals following cluster-based permutation, where applicable. Time 0 s represents picture onset in all plots.

Comparing these power decreases across sessions yielded no significant results, meaning that there were no systematic changes between the group-level context effects from session 1 to session 2.

The underlying sources of the context effect are shown in Figure 5B. Below the specified threshold of *p* < .001, sources were reconstructed exclusively in RH areas, distributed across right inferior frontal gyrus, anterior temporal lobe, inferior parietal lobule and precuneus, lateral occipital cortex, cingulate gyrus, and paracentral lobule.

#### 3.3.3. Picture: Visually guided word retrieval

After picture onset, we evaluated the differences between ERPs of bare and scrambled as well as constrained and unconstrained pictures, shown in Figure 7. For bare versus scrambled, tapping into word retrieval while controlling for visual processing and articulatory preparation, we found significant differences between conditions (orange lines, Figure 7) during the P2 peak window between 250 and 350 ms after picture onset for all groups and sessions except the LH group in session 1 (Control: session 1 ps < .004, session 2 ps < .013; RH: session 1 ps < .032, session 2 ps < .016; LH: session 1 p = .1641, session 2 p = .0391).

**Figure 7.**
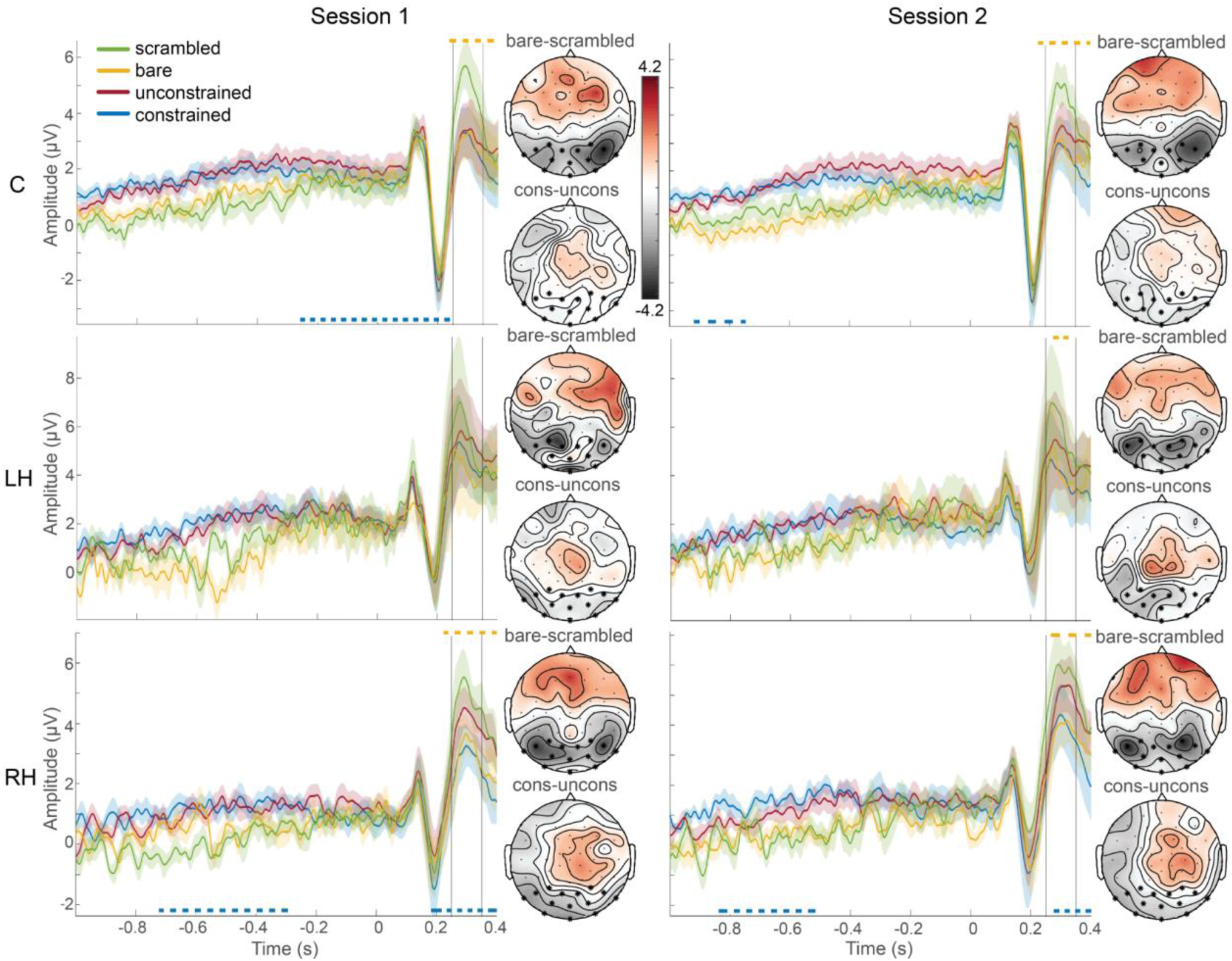
Scalp ERPs from −1000 to 400 ms relative to picture onset for all picture conditions (constrained, unconstrained, bare, scrambled) for 14 occipital electrodes per session (columns, session 1, session 2) and group (rows, C: control, LH: left hemisphere, RH: right hemisphere). The scalp locations of the 14 electrodes are marked in the topographic plots with an asterisk. Color-shaded areas around waveforms represent standard error of the mean. Vertical dashed lines show significant differences between bare and scrambled pictures (orange) and constrained and unconstrained pictures (blue). Differences between conditions during the P2 peak from 250 to 350 ms after picture onset are shown as topographies for constrained minus unconstrained (top) and bare minus scrambled (bottom) per plot, marked by the two vertical lines in each ERP plot. Time 0 s represents picture onset in all plots.

For constrained versus unconstrained pictures, tapping into the lexical-semantic stage of word retrieval, the results revealed significant differences between conditions (blue lines, Figure 7) only in the control and the RH group (Control: session 1 ps < .0377, session 2 p = .0073; RH: session 1 ps < .043, session 2 p = .0371), but not the LH group (session 1 p = .1172; session 2 p = .5781). In the control group, the differences between constrained and unconstrained pictures emerged around picture onset in session 1 (−258 to 240 ms) and sentence offset in session 2 (−918 to −746 ms). For the RH group, they occurred before as well as after picture onset in session 1 (−724 to −294 ms; 182 to 400 ms) and session 2 (−838 to −516; 276 to 400 ms).

While scrambled pictures always evoked the highest P2 peak amplitude, the peak amplitude of the remaining three conditions varied across groups and sessions, but mostly followed the previously established order of unconstrained, bare, and constrained pictures with decreasing peak amplitude. As for the previous two contrasts, we did not detect any systematic amplitude changes between session 1 and 2 within each group when statistically comparing the difference waves for constrained and unconstrained as well as bare and scrambled picture ERPs across sessions.

Sources during the P2 peak from 250 to 350 ms after picture onset for bare versus scrambled and constrained versus unconstrained pictures below *p* < .01 in the control group are shown in Figure 5C and D, respectively. Differences between bare and scrambled pictures were reconstructed in bilateral posterior cortices including precuneus and medial ventral occipital cortex, as well as right paracentral and inferior parietal lobule, and left superior and middle temporal lobe (Figure 5C). For constrained versus unconstrained pictures, sources were reconstructed in the right middle temporal gyrus, inferior parietal lobule, lateral occipital cortex, and sensorimotor regions (postcentral gyrus), as well as the hippocampus and bilateral cerebellar structures (Figure 5D). Note that the control group did not reveal significant differences between constrained and unconstrained pictures during the P2 peak on scalp-level. This could partly explain why the source results for this contrast required a lower threshold than the sentence and the pre-picture results.

#### 3.3.4 Changes in hemispheric dominance

As reported, lesion size did not differ across groups or sessions (3.1, *Lesion Characteristics*). An additional linear model verified that lesion size in session 1 did not systematically co-vary with hemispheric dominance (while controlling for an interaction with group), as this is relevant for interpreting changes in hemispheric dominance between the groups. The overall model was not significant (*F*(3, 14) = 0.16, *p* = .920, *R*² = .034), with lesion size in session 1 not systematically co-varying with hemispheric dominance in session 1 (Estimate = 0.02, *p* = .979).

Changes in hemispheric dominance over time as a function of group are shown stacked across contrasts in Figure 8 (for contrast-specific results, see Supplementary Figure 1). In predicting change in hemispheric dominance between sessions while controlling for contrast type, there was a significant interaction between the lesioned hemisphere (group) and hemispheric dominance in session 1 (Estimate = −1.11, *SE* = 0.48, *z* = −2.30, *p* = .022). In the LH group, about half of the cases were LH-dominant in session 1, and the majority of those changed to right-hemispheric dominance in session 2 (Figure 8, left). At the same time, all but one RH-dominant cases in session 1 remained RH-dominant in session 2. Simple-effects results supported this description, as in the LH group, left hemispheric dominance in session 1 was associated with a change in hemispheric dominance in session 2 (Estimate = −4.07, *SE* = 1.40, *z* = −2.91, *p* = .004). For the RH group (Figure 8, right), simple effects indicated no systematic changes in hemispheric dominance over time (Estimate = 0.37, *SE* = .98, *z* = 0.37, *p* = .709).

**Figure 8.**
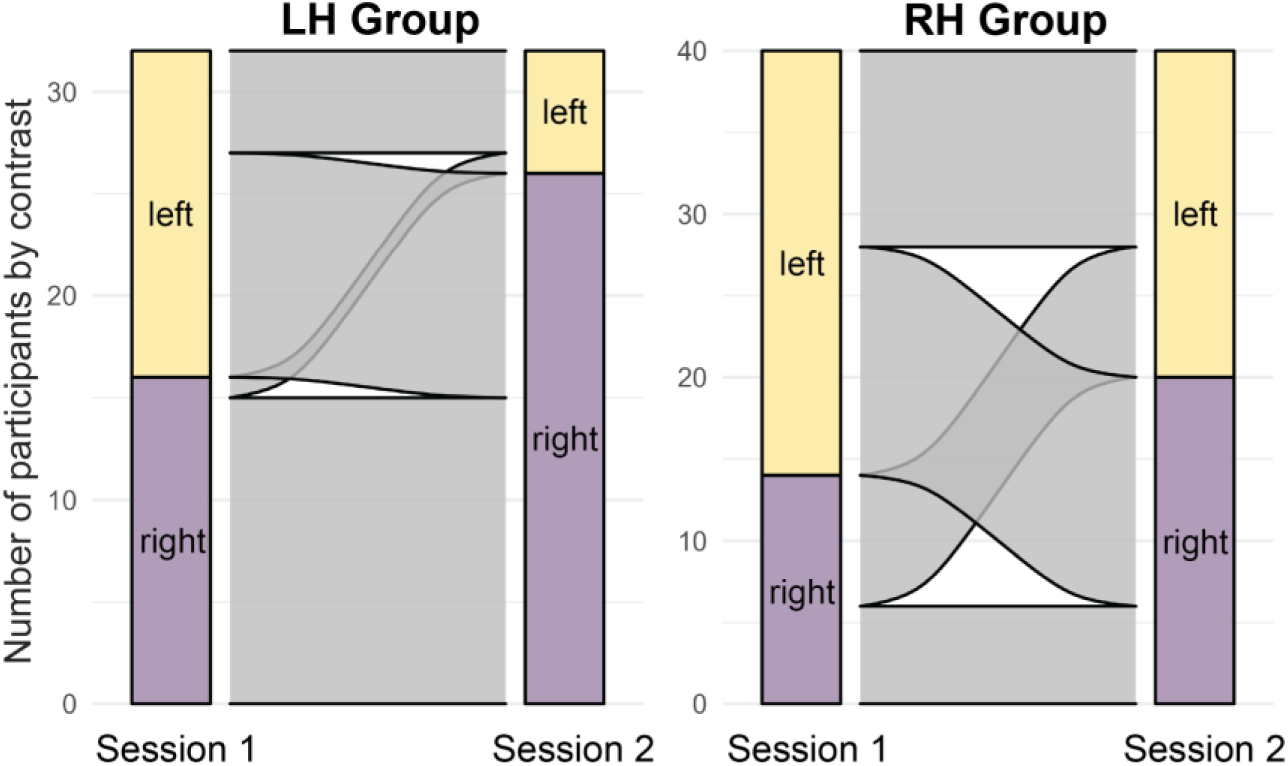
Alluvial plots illustrating changes in hemispheric dominance (left = LH dominant, yellow, right = RH dominant, purple) across all four contrasts (grey flow) for the LH group (left panel) and the RH group (right panel) from session 1 (left column) to session 2 (right column).

## 4. Discussion

We performed a longitudinal study collecting behavioral, EEG, and structural MRI data during the late subacute (2 months) and chronic (7 months) phases of recovery after left and right hemisphere stroke. The inclusion of a stroke-free control group served as a baseline reference for performing state-of-the-art EEG scalp and source-level analyses based on individual MRI scans for the stroke groups. This constituted the study’s main focus of applying state-of-the-art methodology for individualized EEG source reconstruction in the presence of stroke lesions. This approach enabled us to gather evidence on language comprehension as well as production after stroke, for which we employed the Concise Language Paradigm (CLaP, Roos et al., 2024) with EEG.

### 4.1 Behavioral performance

On group-level, the results suggest considerable recovery over time for the LH group, not only in terms of the neuropsychological tests, but also picture naming times and accuracy for the CLaP. Overall, these results are consistent with expected characteristics after left and right hemisphere stroke (Berthier, 2005; Hewetson et al., 2018; Hier et al., 1983), supporting the representativeness of the LH and RH stroke group in our study and increasing confidence in the generalizability of the remaining results.

### 4.2 EEG results in stroke-free controls

Overall, the EEG scalp results of the control group were in line with previous findings for the CLaP from a group of stroke-free control participants (Roos et al., 2024), remaining stable across sessions. For sentence comprehension, the scalp-level results revealed significant amplitude differences between real and reversed speech from 300 to 450 ms after sentence onset, reflecting an earlier decay of the brain signal in response to non-speech input (Stoppelman et al., 2013). On the source-level, these differences manifested in typical left-lateralized language areas in the frontal and temporo-parietal cortex, as well as the right primary auditory cortex. This is in line with previous studies contrasting real and reversed speech using electrocorticography and fMRI (Forseth et al., 2018), or fMRI only (Crinion & Price, 2005).

The context effect between constrained and unconstrained trials during the pre-picture interval on scalp-level yielded alpha-beta power decreases, associated with processes of contextually guided word retrieval (Klaus et al., 2020; Piai et al., 2014, 2015, 2017, 2018; Roos & Piai, 2020). The corresponding source-level results were observed in the right inferior frontal and anterior temporal lobe, and right inferior parietal lobule. This overlaps with areas involved in contextually guided naming of action or object pictures, using similar auditory picture naming tasks with EEG (Chupina et al., 2026; Gastaldon et al., 2020), but differs in laterality from previous MEG results in young adults, for whom this pattern of sources was strongly left-lateralized (Piai et al., 2015; Roos & Piai, 2020).

For visually guided word retrieval, scrambled pictures repeatedly evoked the highest P2 peak on scalp-level, likely reflecting a response to an unexpected visual stimulus inducing a comparably high peak amplitude (for discussion, see Roos et al., 2024). Source-level differences between bare and scrambled pictures during the P2 window manifested in the left temporal as well as the right inferior parietal and occipital lobe. While the occipital sources are in line with electrophysiological processes in response to visual stimuli (Liljeström et al., 2009; Sörös et al., 2003), previous MEG studies also provided evidence for the involvement of right inferior parietal regions in picture naming, linking it to lexical selection processes (Levelt et al., 1998; Liljeström et al., 2009; Sörös et al., 2003). In addition, the left middle and superior temporal lobe have been shown to be involved in lexical-semantic word retrieval and phonetic encoding during spoken word production, respectively (de Zubicaray, 2023; Indefrey, 2011).

Differences between constrained and unconstrained pictures on scalp-level manifested before the P2 component in the control group. This suggests that it derived from differences in sentence context, reflecting processes of contextually guided word retrieval before picture onset in constrained trials, as captured during the pre-picture interval. On the source-level, the differences between constrained and unconstrained pictures during the P2 window occurred over the right occipito-temporal and parietal cortex, including bilateral cerebellar structures. The occipito-parietal areas are similar (but more spread out) to the ones observed for contextually and visually guided naming described above. In fact, the lateral occipital cortex as well as the inferior parietal lobe in the right hemisphere were observed in contextually as well as both visually guided naming contrasts, suggesting an involvement in word-retrieval processes as previously evidenced for its homologous counterpart in the left hemisphere (Roos et al., 2023; Roos & Piai, 2020).

### 4.3 EEG in the stroke groups and changes in hemispheric dominance

Post-stroke EEG results typically show higher variability than control groups. Although the scalp-level results of the stroke groups in our study did not reveal systematic changes in amplitude or power longitudinally (as in the control group), they are not fully in line with the results of the control group. In fact, sentence comprehension was the only contrast where the effects of both stroke groups aligned with the control group by showing a more pronounced ERP peak in response to real speech than reversed speech sentences within the same time window. The remaining contrasts showed more variability.

One could argue that the sentence comprehension contrast in our paradigm, where the effect emerged within the first 450 ms after sentence onset, reflects a more simplistic and cognitively more low-level contrast of speech versus non-speech auditory stimuli, rather than actual speech comprehension throughout the course of a sentence. The pre-picture and picture contrasts, in comparison, might involve more high-level cognitive processes related to word retrieval and picture naming, potentially making them (cognitively) more susceptible to brain damage (and thus potentially more variable). One underlying reason for the divergence between the control and the stroke groups for these contrasts could thus be impacted cognitive processing due to stroke. However, as stroke lesions also affect the spatial distribution and conductivity in the brain, influencing the flow of electrical currents, EEG scalp-level results can be distorted due to stroke as well. By only looking at the scalp-level, we can therefore not determine whether the differences between groups are only cognitive or also trivially physiological in nature.

To add to the interpretation of the scalp results for the stroke groups, we performed individualized source reconstruction, allowing us to control for potential distortions of the stroke lesions at both time points. Using the finite element method (FEM, Vorwerk et al., 2018), we computed detailed three-dimensional models describing the geometry of the different tissues in the head based on session-specific structural MRI scans per participant. By defining mirrored ROIs per contrast based on the session-averaged control group-results, we allowed for changes to homotopic areas in the stroke participants and tailored our analyses to evaluate laterality changes over time, addressing the roles of the left and right hemispheres. Classifying the observed source-level activity in these ROIs into left versus right hemispheric dominance per participant and contrast revealed different results per group. In the LH group, we observed an equal distribution of left versus right hemisphere dominance over contrasts in session 1, followed by systematic changes over time, where the majority of participants for whom the contrasts were left-hemisphere dominant in session 1 changed to right-hemisphere dominance in session 2. The RH group, instead, revealed a rather mixed picture with slightly more left-dominant cases in session 1 and an equal distribution in session 2, with no clear pattern of systematic changes in hemispheric dominance over time.

For the contrasts of interest in our task, this means that LH stroke induced right-hemisphere dominance around seven months post-stroke, if the participant was left-hemisphere dominant for this contrast around two months post-stroke. For LH participants who were already right-hemisphere dominant for that contrast around two months, a change in hemispheric dominance over time was highly unlikely and happened in only one case. Due to the relatively small sample sizes in the current study, we did not explicitly test for changes in hemispheric dominance per contrast type. Of note, all of our contrasts were related to (different aspects of) language processing, and hemispheric dominance within a person can still differ across different aspects of language (Woodhead et al., 2019). Whether and in which direction hemispheric dominance changes could thus differ per contrast within the same participant (not necessarily determined by lesion location alone). It therefore seems highly valuable to employ a paradigm such as the CLaP, which allows to examine the underlying dynamics and individual variation in hemispheric dominance across different contrasts (Bradshaw et al., 2017).

Interestingly, the pattern of changes in hemispheric dominance in the LH group holds despite rather unsystematic underlying lesion locations, which were distributed throughout the left hemisphere without much overlap between participants (Figure 2). At the same time, the lesions of the RH group were more systematic in terms of location and overlap (Figure 2), but did not lead to systematic changes in hemispheric dominance. Overall, we found no systematic changes in lesion size across groups and sessions. Our rationale was based on the framework that neuroplasticity-induced recovery of language tends to happen in the first three months post-stroke (Berthier, 2005; Kauhanen et al., 2000), and that the best behavioral outcome is associated with pre-stroke brain organization (Coleman et al., 2017; Cramer, 2008). In session 1, two-months post-stroke, we thus aimed to capture the state of hemispheric dominance after an initial change in response to the stroke damage, as well as a potential return to pre-stroke brain organization in session 2, seven months post-stroke. However, recovery trajectories can differ individually and we have no information about hemispheric dominance or additional changes thereof before session 1, after session 2, or in between. It is therefore possible that the changes that we captured reflect different stages of recovery across participants.

Nevertheless, these results are based on time-resolved EEG data focusing on specific types of neurophysiological responses and corresponding time-windows of interest within a trial, thus capturing specific processes that are relevant for the employed language task. This suggests that also the changes in hemispheric dominance observed reflect relevant changes in *neural* activity during language use. Specifically, the systematicity of changes towards RH dominance in the LH group, together with the overall improvement of behavioral performance over time, suggests this to be a supportive pattern for recovery in the context of our study. Especially regarding fMRI-based findings of supportive RH-involvement after LH stroke (Saur et al., 2006; Thulborn et al., 1999; Turkeltaub et al., 2026), our results corroborate this pattern by providing complementary EEG source-level evidence. Importantly, these changes were neurophysiological-response specific, adding to the interpretation of the computations performed by the right hemisphere.

Taken together, our study demonstrates that EEG source reconstruction provides a feasible and valid approach to investigate neural activity during post-stroke recovery, provided that the required source methodology is applied including detailed volumetric head models based on individual participant MRI scans per testing session. In contrast to fMRI, which the majority of previous findings are based on, we can be more confident that our findings reflect underlying changes in neural activity occurring *during* the processes themselves, rather than a potentially unreliable abnormality or absence of the BOLD signal after stroke.

### 4.4 Strengths and limitations

In the present study, we used detailed individual FEM forward models accounting for post-stroke changes to volume conduction to perform source reconstruction. Using the CLaP, we were able to evaluate contrasts of language comprehension as well as production during task-relevant time windows (for further strengths and limitations regarding the CLaP specifically, see Roos et al., 2024). Our longitudinal design allowed us to examine changes in hemispheric dominance from two to seven months after stroke in specific ROIs, based on a study-specific stroke-free control group. In addition to a LH stroke group, we also included a RH stroke group, which enabled us to disentangle the impact of LH versus RH stroke on different outcomes, adding to the interpretation of our findings.

Although the source reconstruction was performed in participants’ native space, our analyses for the stroke groups were predetermined by the (mirrored) ROIs resulting from the control group. This enabled us to evaluate hemispheric dominance at both time points. However, we did not relate the change in hemispheric dominance to the original source results of the control group, which also indicated dominance of one hemisphere over the other. In fact, most of the contrasts in the control group indicated a RH bias, which is in line with findings of increased bilateral involvement during aging (Cabeza et al., 2002; Nurmi et al., 2025). Along these lines, future studies should also focus on more individual participant analyses by explicitly taking into account lesion locations and relating these to the observed patterns of hemispheric dominance per participant, as well as to changes in behavioral performance. Since we do not have information about therapy intervention between the late subacute and chronic phase per participant, we cannot disentangle spontaneous recovery from therapy induced improvements.

## 5 Conclusion

We tested participants two and seven months after left or right hemisphere stroke to capture neural activity in the late subacute and chronic stages of post-stroke recovery and evaluate changes in hemispheric dominance between these two time points. The main objective was to employ state-of-the-art EEG scalp and source-level methodology after stroke, for which the obtained results demonstrated the feasibility and validity of our approach. Together, the results provide insights into post-stroke neuroplasticity during language comprehension as well as production, with tentative evidence for a supportive role of the RH *during* language processing at the time scale with which these processes occur.

## Footnotes

1 We note that the term recovery is used inconsistently in the literature, with an ongoing debate as to whether it refers solely to activation of premorbidly involved neural circuits, or also encompasses forms of adaptation through neural plasticity (Code, 2001; Gleeson & Jagoe, 2023). Here, we remain agnostic to this debate and use the term recovery and improvement interchangeably.

## References

Arheix-Parras, S., Glize, B., Guehl, D., & Python, G. (2023). Electrophysiological Changes in Patients with Post-stroke Aphasia: A Systematic Review. Brain Topography, 36(2), 135–171. 10.1007/s10548-023-00941-4

Bates, D., Mächler, M., Bolker, B., & Walker, S. (2015). Fitting Linear Mixed-Effects Models Using lme4. Journal of Statistical Software, 67, 1–48. 10.18637/jss.v067.i01

Berthier, M. L. (2005). Poststroke Aphasia. Drugs & Aging, 22(2), 163–182. 10.2165/00002512-200522020-00006

Blomert, L., Koster, C., & Kean, M.-L. (1995). Antat: Amsterdam-Nijmegen Test voor Alledaagse Taalvaardigheden. Swets & Zeitlinger.

Boersma, P., & Weenink, D. (2017). Praat, software for speech analysis and synthesis. University of Amsterdam.

Bradshaw, A. R., Thompson, P. A., Wilson, A. C., Bishop, D. V. M., & Woodhead, Z. V. J. (2017). Measuring language lateralisation with different language tasks: A systematic review. PeerJ, 5, e3929. 10.7717/peerj.3929

Brodeur, M. B., Dionne-Dostie, E., Montreuil, T., & Lepage, M. (2010). The Bank of Standardized Stimuli (BOSS), a New Set of 480 Normative Photos of Objects to Be Used as Visual Stimuli in Cognitive Research. PLoS ONE, 5(5). 10.1371/journal.pone.0010773

Bullier, B., Cassoudesalle, H., Villain, M., Cogné, M., Mollo, C., De Gabory, I., Dehail, P., Joseph, P.-A., Sibon, I., & Glize, B. (2020). New factors that affect quality of life in patients with aphasia. Annals of Physical and Rehabilitation Medicine, 63(1), 33–37. 10.1016/j.rehab.2019.06.015

Cabeza, R., Anderson, N. D., Locantore, J. K., & McIntosh, A. R. (2002). Aging Gracefully: Compensatory Brain Activity in High-Performing Older Adults. NeuroImage, 17(3), 1394–1402. 10.1006/nimg.2002.1280

Cappa, S. F., Perani, D., Grassi, F., Bressi, F., Alberoni, M., Franceschi, M., Bettinardi, M., Todde, M., & Fazio, M. (1997). A PET Follow-up Study of Recovery after Stroke in Acute Aphasics. Brain and Language, 56(1), 55–67. 10.1006/brln.1997.1737

Chupina, I., Piai, V., & Westner, B. U. (2026). Posterior language areas share electrophysiological signatures of word retrieval in context-driven object and action naming (p. 2026.05.01.721957). bioRxiv. 10.64898/2026.05.01.721957

Chupina, I., Sierpowska, J., Zheng, X. Y., Dewenter, A., Piastra, M.-C., & Piai, V. (2022). Time course of right-hemisphere recruitment during word production following left-hemisphere damage: A single case of young stroke. European Journal of Neuroscience, 56(8), 5235–5259. 10.1111/ejn.15813

Code, C. (2001). Multifactorial Processes in Recovery from Aphasia: Developing the Foundations for a Multileveled Framework. Brain and Language, 77(1), 25–44. 10.1006/brln.2000.2420

Coleman, E. R., Moudgal, R., Lang, K., Hyacinth, H. I., Awosika, O. O., Kissela, B. M., & Feng, W. (2017). Early Rehabilitation After Stroke: A Narrative Review. Current Atherosclerosis Reports, 19(12), 59. 10.1007/s11883-017-0686-6

Cramer, S. C. (2008). Repairing the human brain after stroke: I. Mechanisms of spontaneous recovery. Annals of Neurology, 63(3), 272–287. 10.1002/ana.21393

Crinion, J., & Price, C. J. (2005). Right anterior superior temporal activation predicts auditory sentence comprehension following aphasic stroke. Brain, 128(12), 2858–2871. 10.1093/brain/awh659

Dale, A. M., Liu, A. K., Fischl, B. R., Buckner, R. L., Belliveau, J. W., Lewine, J. D., & Halgren, E. (2000). Dynamic Statistical Parametric Mapping: Combining fMRI and MEG for High-Resolution Imaging of Cortical Activity. Neuron, 26(1), 55–67. 10.1016/S0896-6273(00)81138-1

De Renzi, E., & Vignolo, L. A. (1962). The Token Test: A sensitive test to detect receptive disturbances in aphasics. Brain, 85(4), 665–678. 10.1093/brain/85.4.665

de Zubicaray, G. I. (2023). The Neural Organization of Language Production: Evidence from Neuroimaging and Neuromodulation. In Language Production (1st Edition, p. 46 pages). Routledge. 10.4324/9781003145790-5

Diwakar, M., Tal, O., Liu, T. T., Harrington, D. L., Srinivasan, R., Muzzatti, L., Song, T., Theilmann, R. J., Lee, R. R., & Huang, M.-X. (2011). Accurate reconstruction of temporal correlation for neuronal sources using the enhanced dual-core MEG beamformer. NeuroImage, 56(4), 1918–1928. 10.1016/j.neuroimage.2011.03.042

El Hachioui, H., Lingsma, H. F., Sandt-Koenderman, M. E., Dippel, D. W. J., Koudstaal, P. J., & Visch-Brink, E. G. (2013). Recovery of aphasia after stroke: A 1-year follow-up study. Journal of Neurology, 260(1), 166–171. 10.1007/s00415-012-6607-2

Fan, L., Li, H., Zhuo, J., Zhang, Y., Wang, J., Chen, L., Yang, Z., Chu, C., Xie, S., Laird, A. R., Fox, P. T., Eickhoff, S. B., Yu, C., & Jiang, T. (2016). The Human Brainnetome Atlas: A New Brain Atlas Based on Connectional Architecture. Cerebral Cortex, 26(8), 3508–3526. 10.1093/cercor/bhw157

Fazekas, F., Chawluk, J., Alavi, A., Hurtig, H., & Zimmerman, R. (1987). MR signal abnormalities at 1.5 T in Alzheimer’s dementia and normal aging. American Journal of Roentgenology, 149(2), 351–356. 10.2214/ajr.149.2.351

Forseth, K. J., Kadipasaoglu, C. M., Conner, C. R., Hickok, G., Knight, R. T., & Tandon, N. (2018). A lexical semantic hub for heteromodal naming in middle fusiform gyrus. Brain, 141(7), 2112–2126. 10.1093/brain/awy120

Gastaldon, S., Arcara, G., Navarrete, E., & Peressotti, F. (2020). Commonalities in alpha and beta neural desynchronizations during prediction in language comprehension and production. Cortex, 133, 328–345. 10.1016/j.cortex.2020.09.026

Gleeson, L., & Jagoe, C. (2023). Recovery in the context of post-stroke aphasia: A concept analysis. Advances in Communication and Swallowing, 26(1), 25–35. 10.3233/ACS-220013

Gross, J., Kujala, J., Hämäläinen, M., Timmermann, L., Schnitzler, A., & Salmelin, R. (2001). Dynamic imaging of coherent sources: Studying neural interactions in the human brain. Proceedings of the National Academy of Sciences, 98(2), 694–699. 10.1073/pnas.98.2.694

Hauk, O. (2016). Only time will tell – why temporal information is essential for our neuroscientific understanding of semantics. Psychonomic Bulletin & Review, 23(4), 1072–1079. 10.3758/s13423-015-0873-9

Hewetson, R., Cornwell, P., & Shum, D. (2018). Social participation following right hemisphere stroke: Influence of a cognitive-communication disorder. Aphasiology, 32(2), 164–182. 10.1080/02687038.2017.1315045

Hier, D. B., Mondlock, J., & Caplan, L. R. (1983). Behavioral abnormalities after right hemisphere stroke. Neurology, 33(3), 337–337. 10.1212/WNL.33.3.337

Hilari, K., Byng, S., Lamping, D. L., & Smith, S. C. (2003). Stroke and Aphasia Quality of Life Scale-39 (SAQOL-39). Stroke, 34(8), 1944–1950. 10.1161/01.STR.0000081987.46660.ED

Hillis, A. E., & Heidler, J. (2002). Mechanisms of early aphasia recovery. Aphasiology, 16(9), 885–895. 10.1080/0268703

Hillis, A. E., Kleinman, J. T., Newhart, M., Heidler-Gary, J., Gottesman, R., Barker, P. B., Aldrich, E., Llinas, R., Wityk, R., & Chaudhry, P. (2006). Restoring Cerebral Blood Flow Reveals Neural Regions Critical for Naming. Journal of Neuroscience, 26(31), 8069–8073. 10.1523/JNEUROSCI.2088-06.2006

Homölle, S., & Oostenveld, R. (2019). Using a structured-light 3D scanner to improve EEG source modeling with more accurate electrode positions. Journal of Neuroscience Methods, 326, 108378. 10.1016/j.jneumeth.2019.108378

Hustá, C., Zheng, X., Papoutsi, C., & Piai, V. (2021). Electrophysiological signatures of conceptual and lexical retrieval from semantic memory. Neuropsychologia, 161, 107988. 10.1016/j.neuropsychologia.2021.107988

Indefrey, P. (2011). The Spatial and Temporal Signatures of Word Production Components: A Critical Update. Frontiers in Psychology, 2. https://www.frontiersin.org/articles/10.3389/fpsyg.2011.00255

Jung, T.-P., Makeig, S., Westerfield, M., Townsend, J., Courchesne, E., & Sejnowski, T. J. (2000). Removal of eye activity artifacts from visual event-related potentials in normal and clinical subjects. Clinical Neurophysiology, 111(10), 1745–1758. 10.1016/S1388-2457(00)00386-2

Kaplan, E., Goodglass, H., & Weintraub, S. (1983). Boston naming test. Lea & Febiger.

Kauhanen, M.-L., Korpelainen, J. T., Hiltunen, P., Määttä, R., Mononen, H., Brusin, E., Sotaniemi, K. A., & Myllylä, V. V. (2000). Aphasia, Depression, and Non-Verbal Cognitive Impairment in Ischaemic Stroke. Cerebrovascular Diseases, 10(6), 455–461. 10.1159/000016107

Kessels, R. P. C., van Zandvoort, M. J. E., Postma, A., Kappelle, L. J., & de Haan, E. H. F. (2000). The Corsi Block-Tapping Task: Standardization and Normative Data. Applied Neuropsychology, 7(4), 252–258. 10.1207/S15324826AN0704_8

Kiran, S. (2012). What is the nature of poststroke language recovery and reorganization? ISRN Neurology, 2012. 10.5402/2012/786872

Klaus, J., Schutter, D. J. L. G., & Piai, V. (2020). Transient perturbation of the left temporal cortex evokes plasticity-related reconfiguration of the lexical network. Human Brain Mapping, 41(4), 1061–1071. 10.1002/hbm.24860

Lenth, R. (2023). emmeans: Estimated Marginal Means, aka Least-Squares Means_. *R Package Version 1.8. 5*. https://cir.nii.ac.jp/crid/1370584340724217473

Levelt, W. J. M., Praamstra, P., Meyer, A. S., Helenius, P., & Salmelin, R. (1998). An MEG Study of Picture Naming. Journal of Cognitive Neuroscience, 10(5), 553–567. 10.1162/089892998562960

Liljeström, M., Hultén, A., Parkkonen, L., & Salmelin, R. (2009). Comparing MEG and fMRI views to naming actions and objects. Human Brain Mapping, 30(6), 1845–1856. 10.1002/hbm.20785

Lindauer, U., Dirnagl, U., Füchtemeier, M., Böttiger, C., Offenhauser, N., Leithner, C., & Royl, G. (2010). Pathophysiological Interference with Neurovascular Coupling – When Imaging Based on Hemoglobin Might Go Blind. Frontiers in Neuroenergetics, 2. 10.3389/fnene.2010.00025

Lu, Y., Mao, L., Wang, P., Wang, C., Hartwigsen, G., & Zhang, Y. (2024). Aberrant neural oscillations in poststroke aphasia. Psychophysiology, 61(11), e14655. 10.1111/psyp.14655

Maris, E., & Oostenveld, R. (2007). Nonparametric statistical testing of EEG- and MEG-data. Journal of Neuroscience Methods, 164(1), 177–190. 10.1016/j.jneumeth.2007.03.024

Murzin, V., Fuchs, A., & Kelso, J. A. S. (2013). Detection of Correlated Sources in EEG using Combination of Beamforming and Surface Laplacian Methods. Journal of Neuroscience Methods, 218(1), 96–102. 10.1016/j.jneumeth.2013.05.001

Nurmi, P., Ala-Salomäki, H., Renvall, H., & Liljeström, M. (2025). Magnetoencephalography reveals adaptive neural reorganization maintaining lexical-semantic proficiency in healthy aging (p. 2025.12.19.695466). bioRxiv. 10.64898/2025.12.19.695466

Oostenveld, R., Fries, P., Maris, E., & Schoffelen, J.-M. (2011). FieldTrip: Open Source Software for Advanced Analysis of MEG, EEG, and Invasive Electrophysiological Data. Computational Intelligence and Neuroscience, 2011, 1–9. 10.1155/2011/156869

Piai, V., & Borges, P. (2023). The Electrophysiology of Language Production. In Language Production. Routledge.

Piai, V., & Francken, J. (2026). Explaining what? Time for nuance in the field of neurobiology of language (Hdtg8_v1). PsyArXiv. https://osf.io/preprints/psyarxiv/hdtg8_v1/

Piai, V., Meyer, L., Dronkers, N. F., & Knight, R. T. (2017). Neuroplasticity of language in left-hemisphere stroke: Evidence linking subsecond electrophysiology and structural connections. Human Brain Mapping, 38(6), 3151–3162. 10.1002/hbm.23581

Piai, V., Oostenveld, R., Schoffelen, J. M., & Piastra, M. C. (2024). The impact of CSF-filled cavities on scalp EEG and its implications. Psychophysiology, 61(10), e14624. 10.1111/psyp.14624

Piai, V., Roelofs, A., & Maris, E. (2014). Oscillatory brain responses in spoken word production reflect lexical frequency and sentential constraint. Neuropsychologia, 53, 146–156. 10.1016/j.neuropsychologia.2013.11.014

Piai, V., Roelofs, A., Rommers, J., & Maris, E. (2015). Beta oscillations reflect memory and motor aspects of spoken word production. Human Brain Mapping, 36(7), 2767–2780. 10.1002/hbm.22806

Piai, V., Rommers, J., & Knight, R. T. (2018). Lesion evidence for a critical role of left posterior but not frontal areas in alpha–beta power decreases during context-driven word production. European Journal of Neuroscience, 48(7), 2622–2629. 10.1111/ejn.13695

Piastra, M. C., Oostenveld, R., Schoffelen, J. M., & Piai, V. (2022). Estimating the influence of stroke lesions on MEG source reconstruction. NeuroImage, 260, 119422. 10.1016/j.neuroimage.2022.119422

Price, C. J., & Crinion, J. (2005). The latest on functional imaging studies of aphasic stroke. Current Opinion in Neurology, 18(4), 429. 10.1097/01.wco.0000168081.76859.c1

Puonti, O., Van Leemput, K., Saturnino, G. B., Siebner, H. R., Madsen, K. H., & Thielscher, A. (2020). Accurate and robust whole-head segmentation from magnetic resonance images for individualized head modeling. NeuroImage, 219, 117044. 10.1016/j.neuroimage.2020.117044

R Core Team. (2017). R: A language and environment for statistical computing. https://www.r-project.org/

Roberts, S. M., Bruce, R. M., Hope, T. M. H., Anderson, S., Woodgate, H., Ledingham, K., Leff, A. P., Green, D. W., & Price, C. J. (2026). The Effect of Age on Post-Stroke Language Outcomes. Journal of Aging Research, 2026(1), 7040010. 10.1155/jare/7040010

Roomer, E., Brok, S., Hoogerwerf, A., & Linn, D. (2011). Handleiding Boston Benoemtaak 2011-Een test voor woordvinding. Bachelorthesis, Hogeschool Utrecht, Utrecht.

Roos, N. M., Chauvet, J., & Piai, V. (2024). The Concise Language Paradigm (CLaP), a framework for studying the intersection of comprehension and production: Electrophysiological properties. Brain Structure and Function. 10.1007/s00429-024-02801-8

Roos, N. M., & Piai, V. (2020). Across-session consistency of context-driven language processing: A magnetoencephalography study. European Journal of Neuroscience, 52(5), 3457–3469. 10.1111/ejn.14785

Roos, N. M., Takashima, A., & Piai, V. (2023). Functional neuroanatomy of lexical access in contextually and visually guided spoken word production. Cortex, 159, 254–267. 10.1016/j.cortex.2022.10.014

Rorden, C. (2025). MRIcroGL: Voxel-based visualization for neuroimaging. Nature Methods, 22(8), 1613–1614. 10.1038/s41592-025-02763-7

Rossini, P. M., Altamura, C., Ferretti, A., Vernieri, F., Zappasodi, F., Caulo, M., Pizzella, V., Del Gratta, C., Romani, G.-L., & Tecchio, F. (2004). Does cerebrovascular disease affect the coupling between neuronal activity and local haemodynamics? Brain, 127(1), 99–110. 10.1093/brain/awh012

Ruiter, M. B., Kolk, H. H. J., Rietveld, T. C. M., Dijkstra, N., & Lotgering, E. (2011). Towards a quantitative measure of verbal effectiveness and efficiency in the Amsterdam-Nijmegen Everyday Language Test (ANELT). Aphasiology, 25(8), 961–975. 10.1080/02687038.2011.569892

Ruiter, M. B., Lotgering, E., & Rietveld, A. C. (2016). Adapted ANELT scenarios for ANELT-CU scoring method. *Unpublished Test Stimuli*.

Ruiter, M. B., Otters, M. C., Piai, V., Lotgering, E. A. M., Theunissen, J. E. M. C., & Rietveld, T. C. M. (2023). A transcription-less quantitative analysis of aphasic discourse elicited with an adapted version of the Amsterdam-Nijmegen Everyday Language Test (ANELT). Aphasiology, 37(10), 1556–1575. 10.1080/02687038.2022.2109124

Saur, D., Lange, R., Baumgaertner, A., Schraknepper, V., Willmes, K., Rijntjes, M., & Weiller, C. (2006). Dynamics of language reorganization after stroke. Brain, 129(6), 1371–1384. 10.1093/brain/awl090

Schellekens, M. M. I., Springer, R. C. S., Boot, E. M., Verhoeven, J. I., Ekker, M. S., Van Alebeek, M. E., Brouwers, P. J. A. M., Arntz, R. M., Van Dijk, G. W., Gons, R. A. R., Van Uden, I. W. M., Den Heijer, T., Van Tuijl, J. H., De Laat, K. F., Van Norden, A. G. W., Vermeer, S. E., Van Zagten, M. S. G., Van Oostenbrugge, R. J., Wermer, M. J. H., … Tuladhar, A. M. (2024). Cognitive trajectory in the first year after first-ever ischaemic stroke in young adults: The ODYSSEY study. Journal of Neurology, Neurosurgery & Psychiatry, 95(6), 571–579. 10.1136/jnnp-2023-332104

Schneck, S. M., Levy, D. F., Entrup, J. L., Yen, M., Eriksson, D. K., Casilio, M., Kasdan, A. V., Walljasper, L., Onuscheck, C. F., Davis, L. T., Kirshner, H. S., de Riesthal, M., & Wilson, S. M. (2026). Independent contributions of language activations in left and right temporal cortex to aphasia outcomes after stroke. Imaging Neuroscience, 4, IMAG.a.1225. 10.1162/IMAG.a.1225

Sörös, P., Cornelissen, K., Laine, M., & Salmelin, R. (2003). Naming actions and objects: Cortical dynamics in healthy adults and in an anomic patient with a dissociation in action/object naming. NeuroImage, 19(4), 1787–1801. 10.1016/S1053-8119(03)00217-9

Stojanoski, B., & Cusack, R. (2014). Time to wave good-bye to phase scrambling: Creating controlled scrambled images using diffeomorphic transformations. Journal of Vision, 14(12), 6–6. 10.1167/14.12.6

Stoppelman, N., Harpaz, T., & Ben-Shachar, M. (2013). Do not throw out the baby with the bath water: Choosing an effective baseline for a functional localizer of speech processing. Brain and Behavior, 3(3), 211–222. 10.1002/brb3.129

Thulborn, K. R., Carpenter, P. A., & Just, M. A. (1999). Plasticity of Language-Related Brain Function During Recovery From Stroke. Stroke, 30(4), 749–754. 10.1161/01.STR.30.4.749

Tuladhar, A. M., van Dijk, E., Zwiers, M. P., van Norden, A. G. W., de Laat, K. F., Shumskaya, E., Norris, D. G., & de Leeuw, F.-E. (2016). Structural network connectivity and cognition in cerebral small vessel disease. Human Brain Mapping, 37(1), 300–310. 10.1002/hbm.23032

Turkeltaub, P. E., Martin, K. C., Laks, A. B., & DeMarco, A. T. (2026). Right hemisphere language network plasticity in aphasia. Brain: A Journal of Neurology, 149(4), 1396–1409. 10.1093/brain/awaf308

Tzourio-Mazoyer, N., Landeau, B., Papathanassiou, D., Crivello, F., Etard, O., Delcroix, N., Mazoyer, B., & Joliot, M. (2002). Automated Anatomical Labeling of Activations in SPM Using a Macroscopic Anatomical Parcellation of the MNI MRI Single-Subject Brain. NeuroImage, 15(1), 273–289. 10.1006/nimg.2001.0978

van Casteren, M., & Davis, M. H. (2006). Mix, a program for pseudorandomization. Behavior Research Methods, 38(4), 584–589. 10.3758/BF03193889

van Drongelen, W., Yuchtman, M., Van Veen, B. D., & van Huffelen, A. C. (1996). A spatial filtering technique to detect and localize multiple sources in the brain. Brain Topography, 9(1), 39–49. 10.1007/BF01191641

Van Veen, B. D., Van Drongelen, W., Yuchtman, M., & Suzuki, A. (1997). Localization of brain electrical activity via linearly constrained minimum variance spatial filtering. IEEE Transactions on Biomedical Engineering, 44(9), 867–880. IEEE Transactions on Biomedical Engineering. 10.1109/10.623056

Veldsman, M., Cumming, T., & Brodtmann, A. (2015). Beyond BOLD: Optimizing functional imaging in stroke populations. Human Brain Mapping, 36(4), 1620–1636. 10.1002/hbm.22711

Verhage, F. (1983). Het coderen van het opleidingsniveau voor researchdoeleinden [Educational classification system for research purposes: Revised version]. Academic Hospital Groningen, State University Groningen.

Vorwerk, J., Oostenveld, R., Piastra, M. C., Magyari, L., & Wolters, C. H. (2018). The FieldTrip-SimBio pipeline for EEG forward solutions. BioMedical Engineering OnLine, 17(1), 37. 10.1186/s12938-018-0463-y

Watila, M. m., & Balarabe, S. a. (2015). Factors predicting post-stroke aphasia recovery. Journal of the Neurological Sciences, 352(1), 12–18. 10.1016/j.jns.2015.03.020

Westner, B. U., Dalal, S. S., Gramfort, A., Litvak, V., Mosher, J. C., Oostenveld, R., & Schoffelen, J.-M. (2022). A unified view on beamformers for M/EEG source reconstruction. NeuroImage, 246, 118789. 10.1016/j.neuroimage.2021.118789

Wilson, S. M., Entrup, J. L., Schneck, S. M., Onuscheck, C. F., Levy, D. F., Rahman, M., Willey, E., Casilio, M., Yen, M., Brito, A. C., Kam, W., Davis, L. T., de Riesthal, M., & Kirshner, H. S. (2023). Recovery from aphasia in the first year after stroke. Brain, 146(3), 1021–1039. 10.1093/brain/awac129

Wilson, S. M., & Schneck, S. M. (2021). Neuroplasticity in Post-Stroke Aphasia: A Systematic Review and Meta-Analysis of Functional Imaging Studies of Reorganization of Language Processing. Neurobiology of Language, 2(1), 22–82. 10.1162/nol_a_00025

Woodhead, Z. V. J., Bradshaw, A. R., Wilson, A. C., Thompson, P. A., & Bishop, D. V. M. (2019). Testing the unitary theory of language lateralization using functional transcranial Doppler sonography in adults. Royal Society Open Science, 6(3), 181801. 10.1098/rsos.181801

