## Supplementary for "The Concise Language Paradigm in left and right hemisphere stroke: a longitudinal source-localized EEG study"

**Supplementary Material**

Supplementary Table 1. Group-level error rates in percent (n in parentheses) per session and condition.

| Group | Condition | Session 1 | Session 2 |
|---|---|---|---|
| **Control** | total | 1.3 (39) | 1.3 (41) |
| | bare | 2.4 | 1.8 |
| | constrained | 0.9 | 1.1 |
| | scrambled | 0.3 | 0.3 |
| | unconstrained | 1.4 | 1.8 |
| **LH** | total | 5.9 (111) | 2.9 (55) |
| | bare | 8.6 | 4.4 |
| | constrained | 4.0 | 2.2 |
| | scrambled | 11.2 | 1.7 |
| | unconstrained | 4.2 | 3.2 |
| **RH** | total | 2.0 (47) | 2.4 (57) |
| | bare | 2.9 | 3.3 |
| | constrained | 1.4 | 2.1 |
| | scrambled | 0 | 0.7 |
| | unconstrained | 2.8 | 2.9 |

**Supplementary Table 2.** Group-level mean and SD (in parentheses) picture naming times in seconds per session and condition.

| Group | Condition | Session 1 | Session 2 |
|---|---|---|---|
| **Control** | bare | 0.949 (0.293) | 0.942 (0.264) |
| | constrained | 0.685 (0.210) | 0.685 (0.216) |
| | scrambled | 0.799 (0.213) | 0.783 (0.196) |
| | unconstrained | 0.878 (0.260) | 0.883 (0.245) |
| **LH** | bare | 1.074 (0.385) | 1.018 (0.445) |
| | constrained | 0.813 (0.344) | 0.736 (0.375) |
| | scrambled | 0.914 (0.334) | 0.787 (0.248) |
| | unconstrained | 1.018 (0.370) | 0.951 (0.392) |
| **RH** | bare | 1.064 (0.321) | 1.078 (0.347) |
| | constrained | 0.826 (0.319) | 0.794 (0.320) |
| | scrambled | 0.915 (0.234) | 0.897 (0.232) |
| | unconstrained | 1.049 (0.354) | 1.011 (0.326) |

**Supplementary Table 3.** Within-group RT simple effects between conditions. Results are averaged over levels of sessions. Consecutive contrasts were used, with *p*-values adjusted using the Tukey method for a family of four estimates.

| Group | Contrast | Estimate | SE | z.ratio | p.value |
|---|---|---|---|---|---|
| **Control** | bare-constrained | 263.8 | 20.78 | 12.698 | <.0001 |
| | bare-scrambled | 158.1 | 26.35 | 6 | <.0001 |
| | bare-unconstrained | 67.6 | 20.78 | 3.253 | 0.0063 |
| | constrained-scrambled | -105.8 | 24.3 | -4.352 | 0.0001 |
| | constrained-unconstrained | -196.2 | 8.43 | -23.282 | <.0001 |
| | scrambled-unconstrained | -90.5 | 24.31 | -3.722 | 0.0011 |
| **LH** | bare-constrained | 277.2 | 22.23 | 12.467 | <.0001 |
| | bare-scrambled | 203.2 | 28.35 | 7.166 | <.0001 |
| | bare-unconstrained | 65.8 | 22.24 | 2.958 | 0.0164 |
| | constrained-scrambled | -74 | 26.15 | -2.829 | 0.0241 |
| | constrained-unconstrained | -211.4 | 10.85 | -19.477 | <.0001 |
| | scrambled-unconstrained | -137.4 | 26.16 | -5.25 | <.0001 |
| **RH** | bare-constrained | 263 | 21.47 | 12.249 | <.0001 |
| | bare-scrambled | 168.1 | 27.2 | 6.178 | <.0001 |
| | bare-unconstrained | 42.1 | 21.47 | 1.963 | 0.2023 |
| | constrained-scrambled | -94.9 | 25.09 | -3.783 | 0.0009 |
| | constrained-unconstrained | -220.8 | 9.66 | -22.855 | <.0001 |
| | scrambled-unconstrained | -125.9 | 25.09 | -5.018 | <.0001 |

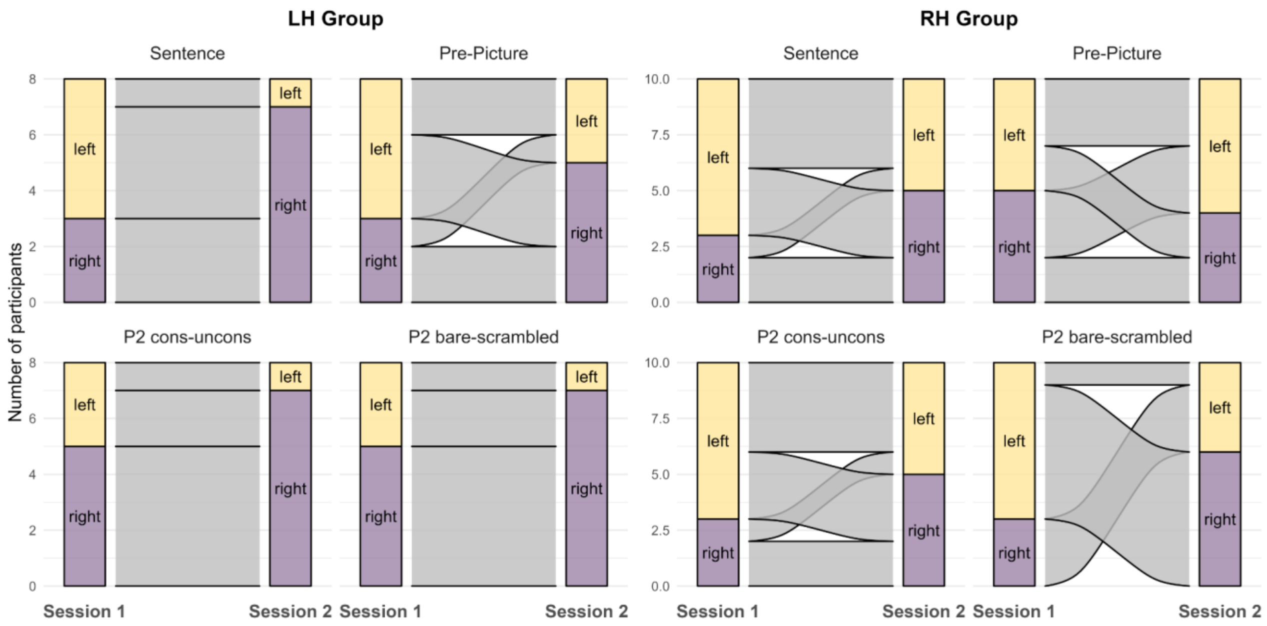


**Supplementary Figure 1.** Alluvial plots illustrating changes in hemispheric dominance (grey flow) for the LH group (left side) and the RH group (right side) from session 1 (left column per panel) to session 2 (right column per panel) per contrast (left = LH dominant, yellow, right = RH dominant, purple).
